# The Nerve Growth Factor IB-like Receptor Nurr1 (NR4A2) Recruits CoREST Transcription Repressor Complexes to Silence HIV Following Proviral Reactivation in Microglial Cells

**DOI:** 10.1101/2021.11.16.468784

**Authors:** Fengchun Ye, David Alvarez-Carbonell, Kien Nguyen, Saba Valadkhan, Konstantin Leskov, Yoelvis Garcia-Mesa, Sheetal Sreeram, Jonathan Karn

**Affiliations:** Department of Molecular Biology and Microbiology. Case Western Reserve University, Cleveland, Ohio, United States of America

**Keywords:** HIV, silencing, microglia, Nurr1, CoREST

## Abstract

Human immune deficiency virus (HIV) infection of microglial cells in the brain leads to chronic neuroinflammation, which is antecedent to the development of HIV-associated neurocognitive disorders (HAND) in the majority of patients. Productively HIV infected microglia release multiple neurotoxins including proinflammatory cytokines and HIV proteins such as envelope glycoprotein (gp120) and transactivator of transcription (Tat). However, powerful counteracting silencing mechanisms in microglial cells result in the rapid shutdown of HIV expression to limit neuronal damage. Here we investigated whether the Nerve Growth Factor IB-like nuclear receptor Nurr1 (NR4A2), which is a repressor of inflammation in the brain, acts to directly restrict HIV expression. HIV silencing was substantially enhanced by Nurr1 agonists in both immortalized human microglial cells (hµglia) and induced pluripotent stem cells (iPSC)-derived human microglial cells (iMG). Overexpression of Nurr1 led to viral suppression, whereas by contrast, knock down (KD) of endogenous Nurr1 blocked HIV silencing. Chromatin immunoprecipitation (ChIP) assays showed that Nurr1 mediates recruitment of the CoREST/HDAC1/G9a/EZH2 transcription repressor complex to HIV promoter resulting in epigenetic silencing of active HIV. Transcriptomic studies demonstrated that in addition to repressing HIV transcription, Nurr1 also downregulated numerous cellular genes involved in inflammation, cell cycle, and metabolism, thus promoting HIV latency and microglial homoeostasis. Thus, Nurr1 plays a pivotal role in modulating the cycles of proviral reactivation by cytokines and potentiating the proviral transcriptional shutdown. These data highlight the therapeutic potential of Nurr1 agonists for inducing HIV silencing and microglial homeostasis and amelioration of the neuroinflammation associated with HAND.

**AUTHOR SUMMARY:** HIV enters the brain almost immediately after infection where it infects perivascular macrophages, microglia and, to a less extent, astrocytes. In previous work using an immortalized human microglial cell model, we observed that integrated HIV constantly underwent cycles of reactivation and subsequent silencing. In the present study, we found that the Nurr1 nuclear receptor is a key mediator of HIV silencing. The functional activation of Nurr1 by specific agonists, or the over expression of Nurr1, resulted in rapid silencing of activated HIV in microglial cells. Global gene expression analysis confirmed that Nurr1 not only repressed HIV expression but also regulated numerous genes involved in microglial homeostasis and inflammation. Thus, Nurr1 is pivotal for HIV silencing and repression of inflammation in the brain and is a promising therapeutic target for treatment of HAND.

## INTRODUCTION

Human immune deficiency virus (HIV) invades the brain soon after primary infection [1]. The virus infects astrocytes, perivascular macrophages, and microglial cells, but not neurons [2, 3]. However, because microglial cells are much longer-lived than astrocytes and perivascular macrophages and can support productive HIV replication, they are mostly likely to be the main cellular reservoir of HIV in the brain [4, 5]. In later stages of HIV infection, many infected patients develop HIV-associated neurocognitive disorders (HAND) [6]. Although combination antiretroviral therapy (cART) dramatically lowers the levels of HIV RNA in the brain [7, 8], it does not reduce the incidence of HAND [9, 10]. Initial studies indicated, paradoxically, that HAND did not correlate with the number of HIV-infected cells or viral antigens in the central nervous system (CNS) [11, 12], but instead correlates strongly with systemic inflammation and CNS inflammation [13]. However, the early studies neglected both the side effects of anti-HIV drugs on neuronal damage, which could mask the benefits of reduced HIV expression by cART and the impact of HIV latency on the development of HAND.

Over the past decade, the intimate relationship between neuroinflammation, neurodegeneration and abnormal activation of microglial cells has been implicated in a wide range of diverse neurological diseases [14–19]. There are compelling reasons to believe that the physiology of microglia also plays a critical role in the development of HAND. Infected macrophages/microglia in the CNS serve as long-lived cellular reservoirs of HIV-1, even in well-suppressed patients receiving ART [20]. Microglia constitute the first barrier of the innate immune response in the brain and become activated and polarized to maintain the integrity of the CNS [21, 22]. In the normal CNS environment, healthy neurons provide signals to microglia via secreted and membrane bound factors such as CX3CL1 and neurotransmitters that induce HIV-silencing. By contrast, damaged neurons not only cause activation of the microglia but also induce HIV reactivation [23].

Activated microglia secrete exaggerated amounts of neurotoxins such as tumor necrosis factor-alpha (TNF-α), nitric oxide, interleukin-6 (IL-6), interleukin-1 beta (IL-1β), reactive free radicals, and matrix metallopeptidases (MMPs) [24, 25]. The production of these cytotoxic factors is augmented by HIV infection [23, 26]. Mounting evidence indicates that HIV proteins such as transactivator of transcription (Tat), negative regulatory factor (Nef), envelope glycoprotein gp120, and viral RNA are not only directly neurotoxic, but also contribute to inflammation in the brain by activating microglial cells [27–35]. On the other hand, some of the inflammatory cytokines such as TNF-α strongly induce HIV expression in microglial cells through autocrine signaling, creating cycles of HIV reactivation and chronic inflammation in the brain [36]. It is therefore important to determine the factors responsible for inducing HIV reactivation and inflammation and explore cellular mechanisms that antagonize these factors in order to develop treatment for HAND.

A major constraint for studying HIV infection and replication in the brain is the difficulty of obtaining native microglial cells from brain biopsies. We therefore developed a microglial cell model by immortalizing human primary microglial cells with the simian virus large T-antigen (SV40) and the human telomerase reverse transcriptase (hTERT) [37]. The immortalized microglia retain the typical structure and morphology of primary microglial cells, express microglial cell markers, and display microglial cell activities such as migration and phagocytosis [37].

A unique feature of HIV infection of microglial cells is that the virus is able to quickly establish latency [23, 36–38]. In microglial cells, transcription initiation is primarily regulated by NF-κB. In resting microglia (M0 stage), NF-κB is sequestered cytoplasm [23, 36–38]. However, unlike memory T-cells, P-TEFb is not disrupted, although inhibited by CTIP2 [39, 40]. The provirus is also silenced epigenetically through the CoREST and polycomb repressive complex 2 (PRC2) histone methyltransferase machinery [4, 41–44]. Activation of microglia by pro-inflammatory signals, such as TNF-α, reversed these molecular restrictions and leads to the reactivation of dormant proviruses and neuropathology.

In contrast to T cells where integrated HIV eventually establishes permanent latency until it is activated by cellular signaling events, HIV in microglial cells undergoes cycles of spontaneous reactivation and subsequent silencing [36]. For example, using a co-culture of (iPSC)-derived human microglial cells (iMG) that were infected with HIV and neurons, we demonstrated that HIV expression in iMG was repressed when co-cultured with healthy neurons but induced when co-cultured with damaged neurons [23]. The dynamics of spontaneous reactivation of latent HIV and subsequent silencing of active HIV constantly typically generates two populations in culture: the GFP^-^ population with transient latent HIV, and the GFP^+^ population undergoing active HIV transcription. Spontaneous HIV reactivation in microglial cells could be attenuated by activation of the glucocorticoid receptor (GR) with its ligand dexamethasone (DEXA) [45], which blocked recruitment of NF-κB and AP-1 for HIV transactivation [36, 45]. However, since we consistently observed spontaneous reactivation of latent HIV and subsequent silencing of the active HIV in the absence of dexamethasone, and in co-cultures with neurons, we reasoned that there exist other cellular factors that promote HIV silencing in microglial cells.

In the present study, we examined whether three members of the Nerve Growth Factor IB-like nuclear receptor family, which includes nuclear receptor 77 (Nur77, NR4A1), nuclear receptor related 1 (Nurr1, NR4A2), and neuron-derived receptor 1 (Nor1, NR4A3), contribute to HIV silencing in microglial cells. These receptors play complementary roles in neurons and microglia to limit inflammatory responses. In neurons, these receptors act as positive transcriptional regulators that control expression of dopamine transporter and tyrosine hydroxylase for differentiation of dopamine neuron, as well as other key genes involved in neuronal survival and brain development [46–49]. By contrast, these nuclear receptors can also act as negative transcriptional regulators in microglia cells and suppress expression of inflammatory cytokines such as TNF-α and IL-1β [50]. Because of these combined mechanisms, Nerve Growth Factor IB-like nuclear receptors play a critical role in protection of the brain during neurodegenerative diseases such as Parkinson’s disease and Alzheimer’s disease [51–56].

Here we report that Nurr1 plays a pivotal role in silencing active HIV in microglial cells by recruiting the CoREST/HDAC1/G9a/EZH2 transcription repressor complex to HIV promoter. Our data also demonstrate that Nurr1 promotes microglial homoeostasis and suppression of inflammation in the brain.

## RESULTS

### Nurr1 agonists strongly induce HIV silencing in microglial cells

To study the role of nuclear receptors in the control of HIV expression in the microglia, we used our immortalized human microglial (hµglia) cells [37], which were infected with a recombinant HIV-1 reporter that carried an EGFP marker for “real-time” monitoring of HIV latency and reactivation (**Fig 1A**). One representative clone, HC69 [37, 45], was used for all experiments described in this study. Under normal culture conditions, most cells were GFP-negative (GFP^-^) (**Fig 1B** & **C**). Exposure of HC69 cells to TNF-α (400 pg/ml) for 24 hours (hr), induced GFP expression (GFP^+^) in over 90% of the cells, demonstrating that majority of the integrated HIV provirus was in a latent state under normal culture conditions. To examine whether the reactivated HIV could revert to latency, we conducted a chase experiment by culturing the activated HC69 cells for 96 hr in fresh medium following TNF-α stimulation for 24 hr and washing with PBS. Notably, the numbers of GFP+ cells decreased from 93.1% to 61.4% at the end of the chase experiment, suggesting the existence of an intrinsic cellular mechanism that silences the activated HIV. This substantial decrease of GFP+ expression was unlikely to be caused by GR-mediated HIV silencing [45], because the cells were cultured in the absence of GR ligand glucocorticoid or dexamethasone.

**Figure 1.**
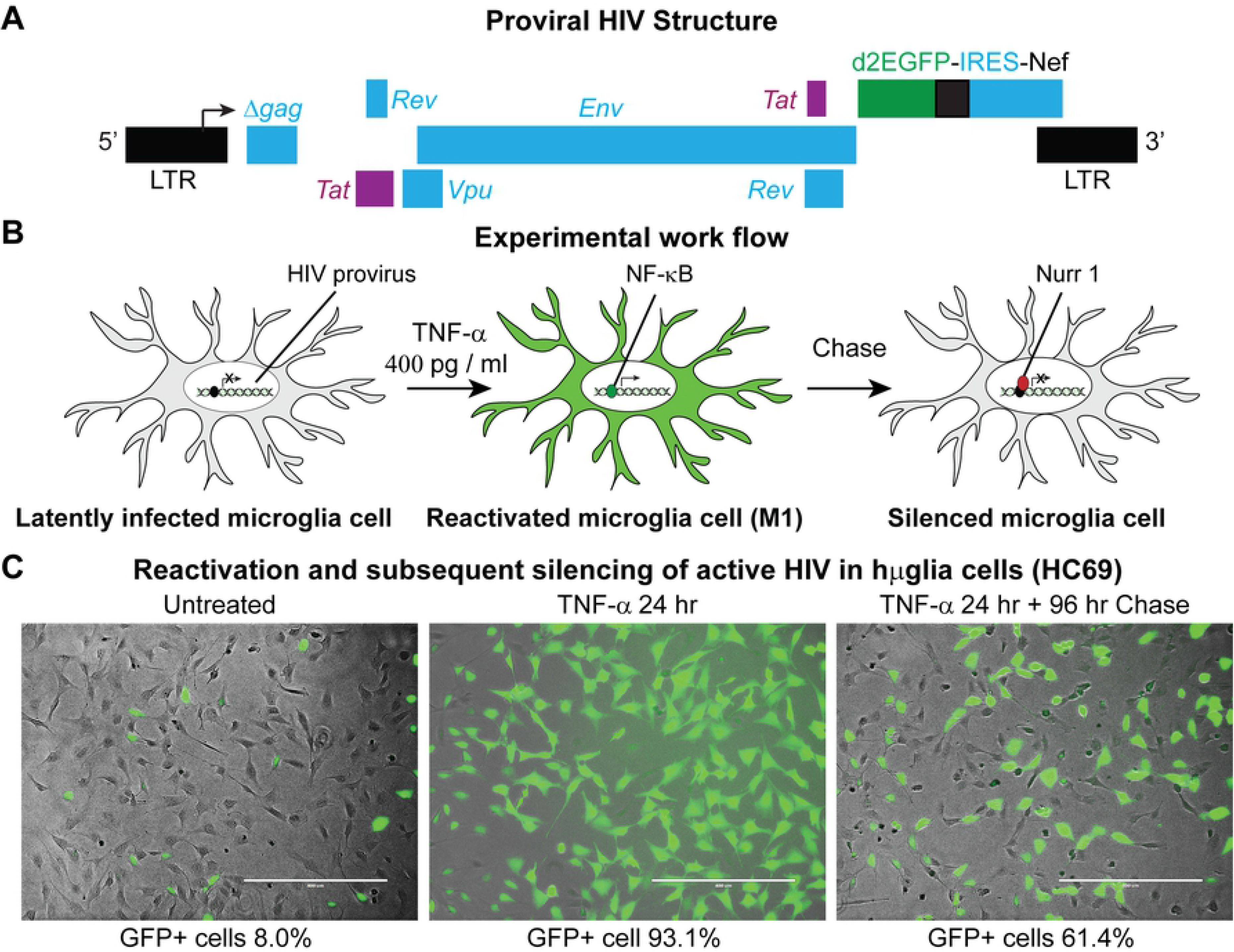
Spontaneous silencing of active HIV in microglial cells. **A**, genome organization of a d2EGFP reporter HIV-1 that was cloned in the lentiviral vector pHR’. A fragment of HIV-1_pNL4-3_, containing *Tat*, *Rev*, *Env*, *Vpu* and *Nef* with the green fluorescence reporter gene d2EGFP, was cloned into the lentiviral vector pHR’. The resulted plasmid was used to produce the VSV-G HIV particles as described previously [120]. Immortalized human microglial cells (hµglia) were infected with the lenti-HIV viral particles, generating multiple clones with an integrated pro-virus genome. HC69 was a representative of these clones. **B,** Schematic diagram of experimental scheme to study the role of nuclear receptors in microglial reactivation and reversion to latency. **C,** Representative phase contrast, GFP, and overlapped images of HC69 cells that were cultured in the absence (untreated, left panel) and in the presence of TNF-α (400 pg/ml) for 24 hr (TNF-α 24 h, middle panel) respectively, or used in a chase experiment by continuously culturing HC69 cells in the absence of TNF-α for 96 hr after stimulating the cells with TNF-α (400 pg/ml) for 24 hr and washing with PBS (TNF-α 24 h+96 h, right panel). The average percentages of GFP+ cells indicated for each panel were measured by flow cytometry from triplicate wells.

To understand the regulatory mechanisms of HIV expression in microglial cells, we had previously undertaken a global screening for HIV silencing cellular factors [57, 58] by using a HIV-infected rat microglial cell model (CHME cells) [37, 59]. The latently infected CHME/HIV cells were superinfected with lentiviral vectors carrying a synthetic shRNA library from Cellecta Inc. (Mountain View, CA) containing a total of 82,500 shRNAs targeting 15,439 mRNA sequences [60–62].. Cells carrying reactivated proviruses were then purified by sorting and the shRNA sequences were identified by next-generation sequencing and classified by Ingenuity Pathway Analysis (QIAGEN). This powerful new technology, which we have also applied to the identification of latency factors in T-cells and TB-infected myeloid cells [58, 63], has revealed a wide range of factors and pathways critical for maintaining proviral latency in microglial cells. Analysis of the top 25 % “hits” led to our unexpected discovery that members of the nuclear receptors (NRs) families including Thyroid Hormone Receptor-like family members PPAR`α`, PPAR`β`, PPAR`γ`, and RAR`β` ranked in the top 5%, the Retinoid X Receptor-like family members RXR`α` and RXR`β` together with the glucocorticoid receptor (GR, NR3C) ranked in the top 15%, and the Nerve Growth Factor IB-like family members NR4A1 (Nur77), NR4A2 (Nurr1) and NR4A3 (Nor1) ranked in the top 25%.

Agonists of the NR4A nuclear receptor family (Nur77 (NR4A1), Nurr1 (NR4A2), and Nor1 (NR4A3)) have been shown to ameliorate neuron degeneration in animal models [53, 64–67]. To confirm a role for the nuclear receptors in HIV silencing, we first treated spontaneously activated HC69 cells with the Nurr1 agonist 6*-*mercaptopurine (6-MP) [68, 69]. As shown in **Fig 2A**, the frequency of GFP^+^ cells decreased in a 6-MP dose-dependent manner. Data from Western blot analysis showed that HC69 cells constitutively expressed Nurr1, as well as a very low level of Nor1 (**Fig 2B**), but Nur77 expression in these cells was below the detection limit. Treatment with 6-MP slightly increased expression of Nurr1. Expression of HIV, as measured by the levels of Nef protein, was strongly inhibited in a dose-dependent manner. Notably, as a control for the role of Nurr1 in cellular gene expression, 6-MP also substantially reduced expression of MMP2, which is a well-known repression target of Nurr1 and a neurotoxin involved in the development of HAND [70, 71].

**Figure 2.**
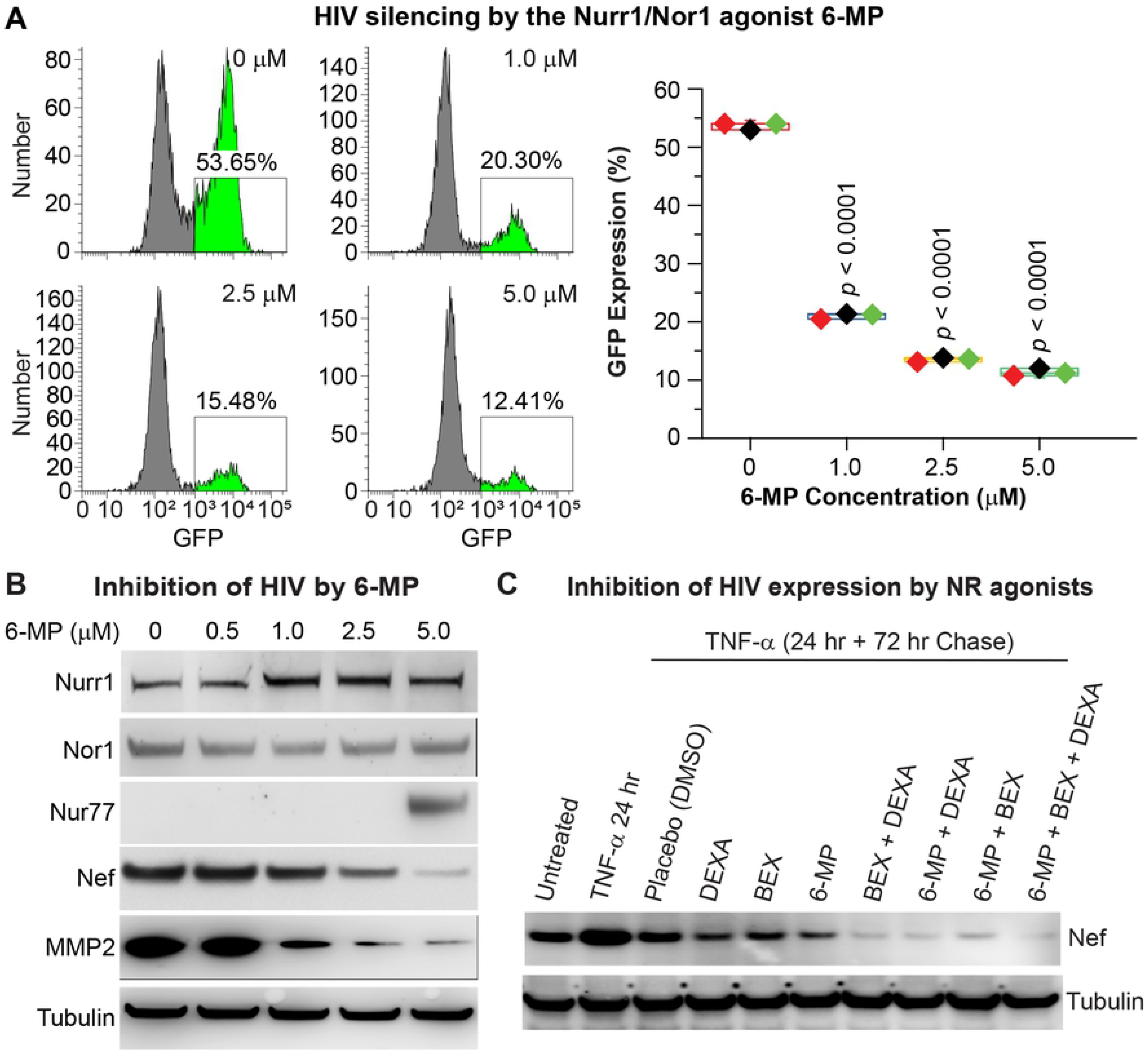
Activation of Nurr1 enhances HIV silencing in immortalized human microglial cells (hµglia). **A**, Impact of Nurr1 agonist 6-MP on HIV expression. Left: Representative flow cytometry histograms. Right: Quantitative results from three independent experiments. For this experiment, we used a batch of HC69 cells with high numbers of GFP^+^ cells resulting from spontaneous HIV reactivation following multiple passages of culture in the absence of dexamethasone. These cells were cultured in the presence of different doses of 6-MP for three days. The percentages of GFP^+^ cells from the differently treated cells were measured by flow cytometry. The *p*-values of pair-sample, Student’s *t*-tests comparing un-treated cells and cells treated with different doses of 6-MP were calculated from three independent experiments. **B**, Western blot detection of Nurr1, Nor1, HIV-1 Nef protein, and Nurr1 target gene MMP2 in HC69 cells described in A. The level of β-tubulin was used as a loading control. **C**, the nuclear receptor agonists dexamethasone (DEXA, 1 µM), Bexarotene (BEX, 1 µM) and 6-MP (1 µM) have additive effects on HIV silencing in HC69 cells. HC69 cells were first treated with high dose (400 pg/ml) TNF-α for 24 hr, followed by a 72 hr chase experiment during which the cells were washed with PBS and cultured in fresh media in the presence of placebo (DMSO) or the various NR agonists, alone or in combination. Expression of Nef and β-tubulin in the differently treated cells was analyzed by Western blot analysis as described in C.

In addition, the Retinoid X Receptor-like family members also play a critical role in silencing inflammation in the brain [72, 73]. We also screened various agonists of the nuclear receptors for their effect on HIV expression in HC69 microglia (**Fig 2C**). We induced maximum HIV expression in HC69 cells with high dose (400 pg/ml) TNF-α for 24 hr, followed by a chase experiment during which the induced cells were cultured in the absence or presence of various agonists, alone or in combination. Consistent with our previous gene manipulation data, the RXRα/β/γ agonist bexarotene (BEX) [74–77] silenced HIV expression on its own, although it was less potent than 6-MP. Interestingly, combinations of 6-MP with DEXA and BEX displayed additive HIV silencing effects, suggesting that they each had distinct mechanisms of action.

### Nurr1 overexpression enhances HIV silencing

To further examine how the nuclear receptors contribute to HIV silencing, we constructed lentiviral vectors expressing N-terminal 3X-FLAG-tagged Nur77, Nurr1, and Nor1 respectively under the control of a CMV promoter. Infection of HC69 cells with the different lentiviruses generated cell lines that stably expressed FLAG-tagged Nur77, Nurr1, Nor1, and empty vector, respectively, as confirmed by RNA-Seq studies (**Fig 3A**) western blots (**Fig 3B**).

**Figure 3.**
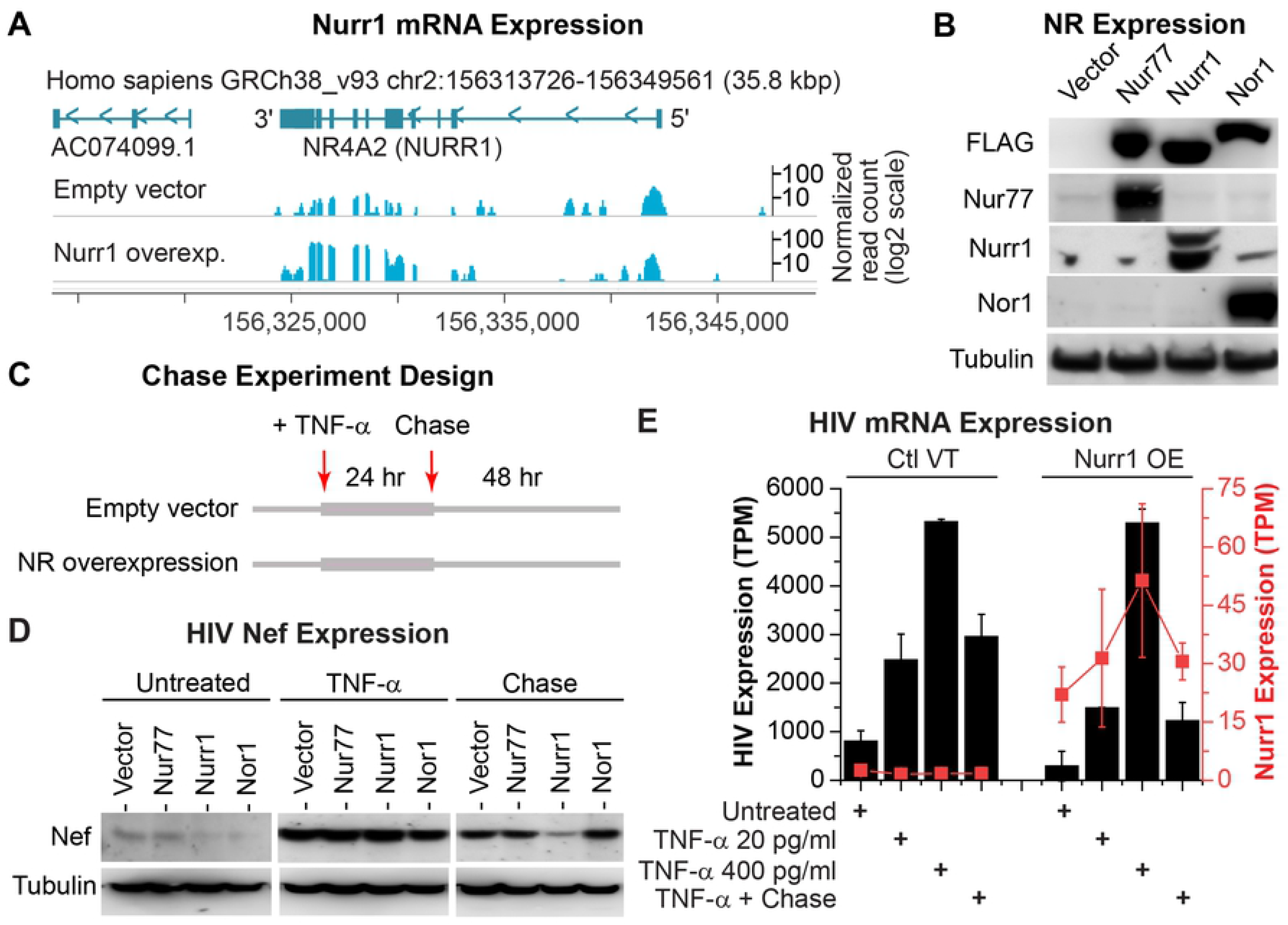
Overexpression of Nurr1 in HC69 cells enhances HIV silencing. **A,** RNA-Seq confirmation of overexpression (OE) of Nurr1 in HC69 cells. Sequence read histograms for the Nurr1 locus is shown for control (vector) and Nurr1 overexpression. Annotated genes for the shown locus are indicated on the top, and the position of the locus on chromosome 2 is shown both at the top and the bottom. A read scale for each row is shown on the right, with the values for the overexpression studies drawn on a log2 scale. **B,** Verification of Nur77, Nurr1, and Nor1 overexpression by Western blot analysis in HC69 cell lines stably expressing 3X-FLAG-tagged Nur77, Nurr1, and Nor1 respectively. HC69 cells stably carrying the 3X-FLAG-empty vector were used as a reference for comparison. The level of β-tubulin was used as a loading control. Notably, the levels of endogenous Nur77 and Nor1 in HC69 cells were very limited. In contrast, Nurr1 was constitutively expressed in HC69 cells. **C,** Schematic depicting the TNF-α stimulation and chase studies. The four cell lines described in B were either untreated or treated with high dose (400 pg/ml) TNF-α for 24 h. To examine HIV silencing, one set of TNF-α induced cells were used in a chase experiment by continuous culture of the cells in the absence of TNF-α for an additional 48 h. The time points at which TNF-α is added or removed are shown by arrows on the top. **D,** Expression of HIV Nef protein in the different cell lines before and after TNF-α stimulation and at the end of the chase experiment was measured by Western blot analysis. The level of β-tubulin was used as a loading control. **E,** Expression level of HIV mRNA (black bar graph) and Nurr1 (red rectangles and lines) in transcripts per million cellular transcripts are shown for each of the treatment steps shown in panel C in both vector-infected cells (on the left) and Nurr1 overexpressing cells (on the right half of the graph). For the 24 hr TNF-α stimulation step, both a low dose (20 pg/ml) and a high dose (400 pg/ml) are used. The values shown are the average of three replicate RNA-Seq samples with two standard deviations as error bars. The expression values for HIV and Nurr1 are shown on Y axes to the left and right, respectively.

To examine how overexpression of each of these nuclear receptors modulated HIV proviral activation and silencing, we stimulated all four cell lines with high dose (400 pg/ml) TNF-α for 24 hr to induce HIV transcription through activation of NF-κB [38], followed by a 48 hr chase experiment in which TNF-α was removed by washing the cells with PBS followed by the addition of media lacking TNF-α (**Fig 3C**). As shown by western blot in **Fig 3D**, TNF-α strongly induced the expression of HIV Nef protein, which we used as a marker of HIV reactivation, in all cell lines at 24 hr. Notably, Nef expression decreased in all four cell lines 48 hr after TNF-α withdrawal. However, the reduction in Nef expression was much more pronounced in HC69 cells that express 3X-FLAG-Nurr1, suggesting that overexpression of Nurr1 enhanced silencing of active HIV in HC69 cells.

We rigorously confirmed these findings using the RNA-Seq data (**Fig 3E**) to measure the fluctuations in both HIV and Nurr1 expression. In the Nurr1 overexpressing cells, even in unstimulated conditions, the level of HIV proviral expression was strongly reduced. Following stimulation with either a low dose (20 pg/ml) or high dose (400 pg/ml) TNF-α, both vector-infected and Nurr1 overexpressing cells showed an increase in proviral expression. While the level of HIV expression was similar between control cells (vector-infected) and Nurr1-overexpressing cells after high dose TNF-α stimulation, Nurr1 overexpressing cells had much lower proviral expression level after low dose TNF-α stimulation (**Fig 3E**). The level of HIV mRNA after withdrawal of high dose TNF-α was three times lower in Nurr1 overexpressing cells than in vector-infected cells (**Fig 3E**), strongly suggesting that overexpression of Nurr1 enhanced silencing of active HIV in HC69 cells.

### Nurr1 knockdown blocks HIV silencing

As a complementary approach we performed shRNA-mediated knock down (KD) of endogenous Nurr1 in HC69 cells. Cell lines that stably expressed Nurr1-specific or control shRNA were verified for effective Nurr1 KD by RNA-Seq analyses (**Fig 4A**) Following the protocol described in **Fig 4B**, control and KD cells were activated with a high dose (400 pg/ml) TNF-α for 24 hr, followed by a 72 hr chase. Western blot analyses confirmed the Nurr1 knock down efficiency (**Fig 4C**). The blots also showed that HIV Nef protein, which is a measure of HIV transcription, was strongly induced at 24 hr post TNF-α stimulation in both the control and the Nurr1 KD cells. However, after the chase, Nef levels decreased significantly in the control cells but remained high in Nurr1 KD cells (**Fig 4C**).

**Figure 4.**
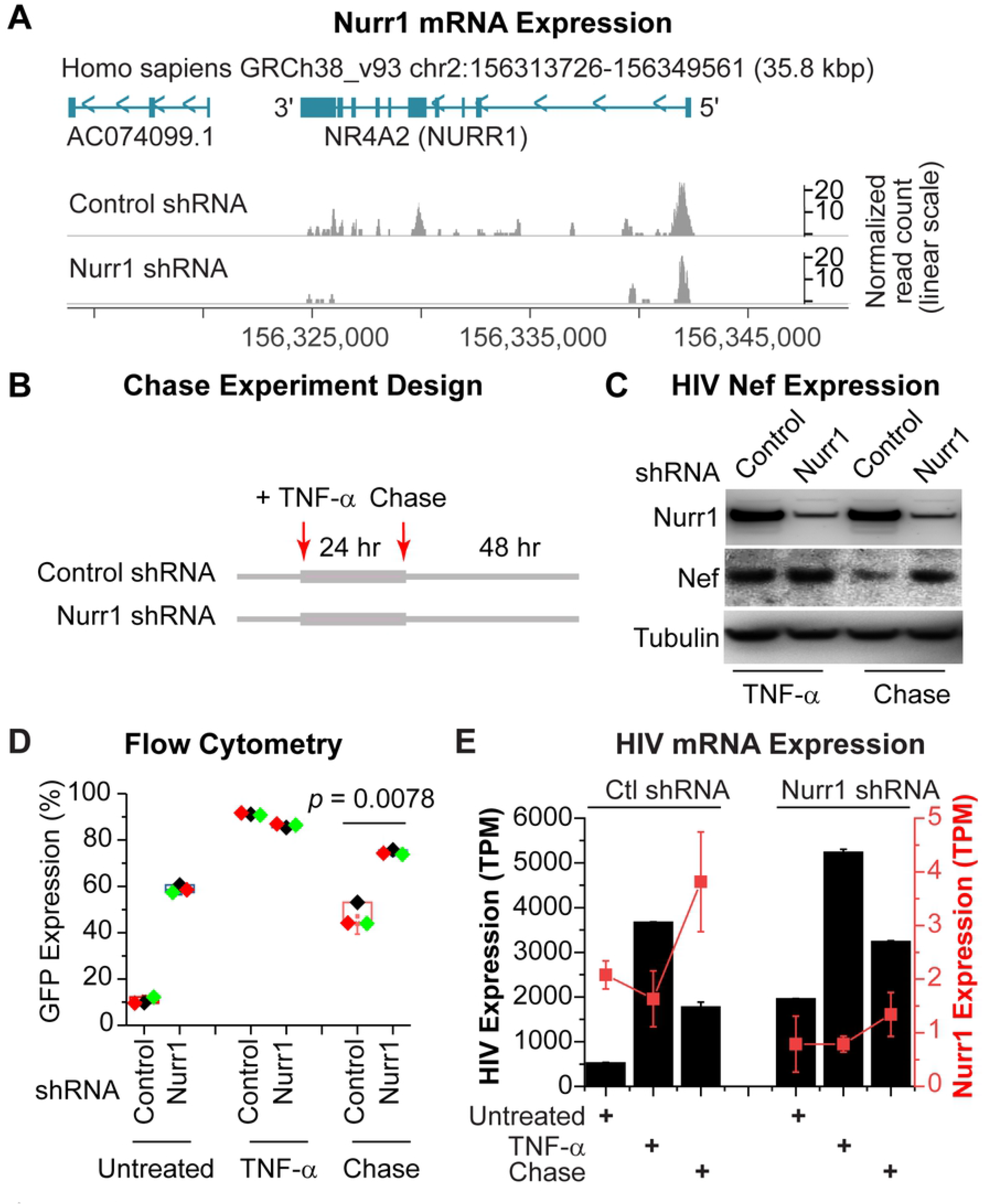
Nurr1 knock down (KD) in HC69 cells enhances HIV expression and block proviral silencing during the chase step. **A,** RNA-Seq confirmation of Nurr1 KD in HC69 cells. Read histograms for the Nurr1 locus is shown for non-targeting shRNA-infected cells, and cells infected with Nurr1 specific shRNA lentiviral constructs. Annotated genes for the shown locus are indicated on the top, and the position of the locus on chromosome 2 is shown both at the top and the bottom. A read scale for each row is shown on the right, with the values for the knock down studies drawn on a linear scale. **B,** Schematic depicting the TNF-α stimulation and chase studies. The two shRNA lentiviral transduced cell lines described in A were either untreated or treated with high dose (400 pg/ml) TNF-α for 24 hr. One set of TNF-α induced cells were used in a chase experiment in the absence of TNF-α for an additional 48 hr. The time points at which TNF-α is added or removed are shown by arrows on the top. **C,** Western blot studies measuring the expression of endogenous Nurr1, Nef, and β-tubulin in cells infected with either a non-targeting control shRNA or Nurr1-specific shRNA lentiviral vectors. The expression patterns from the TNF-α (400 pg/ml) stimulation and the chase step are shown. **D,** KD of endogenous Nurr1 strongly inhibits HIV silencing. The percentages of GFP^+^ cells in the two cell lines, before treatment, at 24 hr post-TNF-α (400 pg/ml) stimulation, and at 72 hr after TNF-α withdrawal (chase) were analyzed by flow cytometry and calculated from three independent experiments. The difference in GFP expression between the two cell lines at 72 hr chase was statistically significant, with a *p* = 0.0078. **E,** Expression level of Nurr1 (red rectangles and lines) and the HIV provirus (black bar graph) in transcripts per million cellular transcripts are shown for each of the treatment steps in both non-targeting shRNA infected cells (on the left) and Nurr1-specific shRNA-infected cells (on the right half of the graph). The values shown reflect the average of three replicate RNA-Seq samples from two distinct shRNA constructs per control and Nurr1 knock down groups, with two standard deviations as error bars. The expression values for HIV and Nurr1 are shown on Y axes to the left and right, respectively.

Similar results were obtained using flow cytometry (**Fig 4D**). Compared to cells expressing control shRNA with 10.5% GFP+ cells, the Nurr1 KD cells displayed 58.8% GFP^+^ cells even before TNF-α stimulation, which most likely resulted from failure of silencing spontaneously reactivated HIV in these cells due to Nurr1 depletion (**Fig 4D**). As expected, after exposure to high dose TNF-α for 24 hr, both the control and Nurr1 KD cell lines expressed equally high levels of GFP expression (**Fig 4D**), displaying 86.3% and 91.2% GFP+ cells respectively. However, 72 hrs after TNF-α withdrawal, GFP expression decreased significantly in cells expressing the control shRNA (47.2% GFP+) but remained high (74.6% GFP+) in the Nurr1 KD cells (**Fig 4D**). Finally, the overall mRNA level of the HIV measured by RNA-Seq was about 1.7 times higher in Nurr1 KD at the end of the chase experiment (**Fig 4E**).

Thus, both the overexpression and the reciprocal KD experiments confirmed an essential role of Nurr1 in the silencing HIV in microglial cells.

### Nurr1 drives activated microglial cells towards homeostasis

Our RNA-Seq data also provided important insights into the cellular pathways that were impacted by Nurr-1 over- and under-expression. We focused our attention on the changes in cellular transcriptome during the chase step following TNF-α induction since, as described above, this is the stage where Nurr1 has the greatest impact on HIV gene expression. As shown by the differential gene expression curves in **Fig 5A,** a small subset of genes are selectively up and down regulated during the chase. A larger number of genes were differentially expressed in Nurr1 overexpressing cells compared to control cells (**Fig 5A & S1 Fig.**). Pathways that showed the most statistically significant changes in response to Nurr1 overexpression included the downregulation of key pathways with critical roles in cellular proliferation and metabolism including: MYC, E2F and MTORC signaling and G2M checkpoint (**Fig 5B**). By contrast, KD of Nurr1 by shRNA did not selectively activate any major signaling pathways.

**Figure 5.**
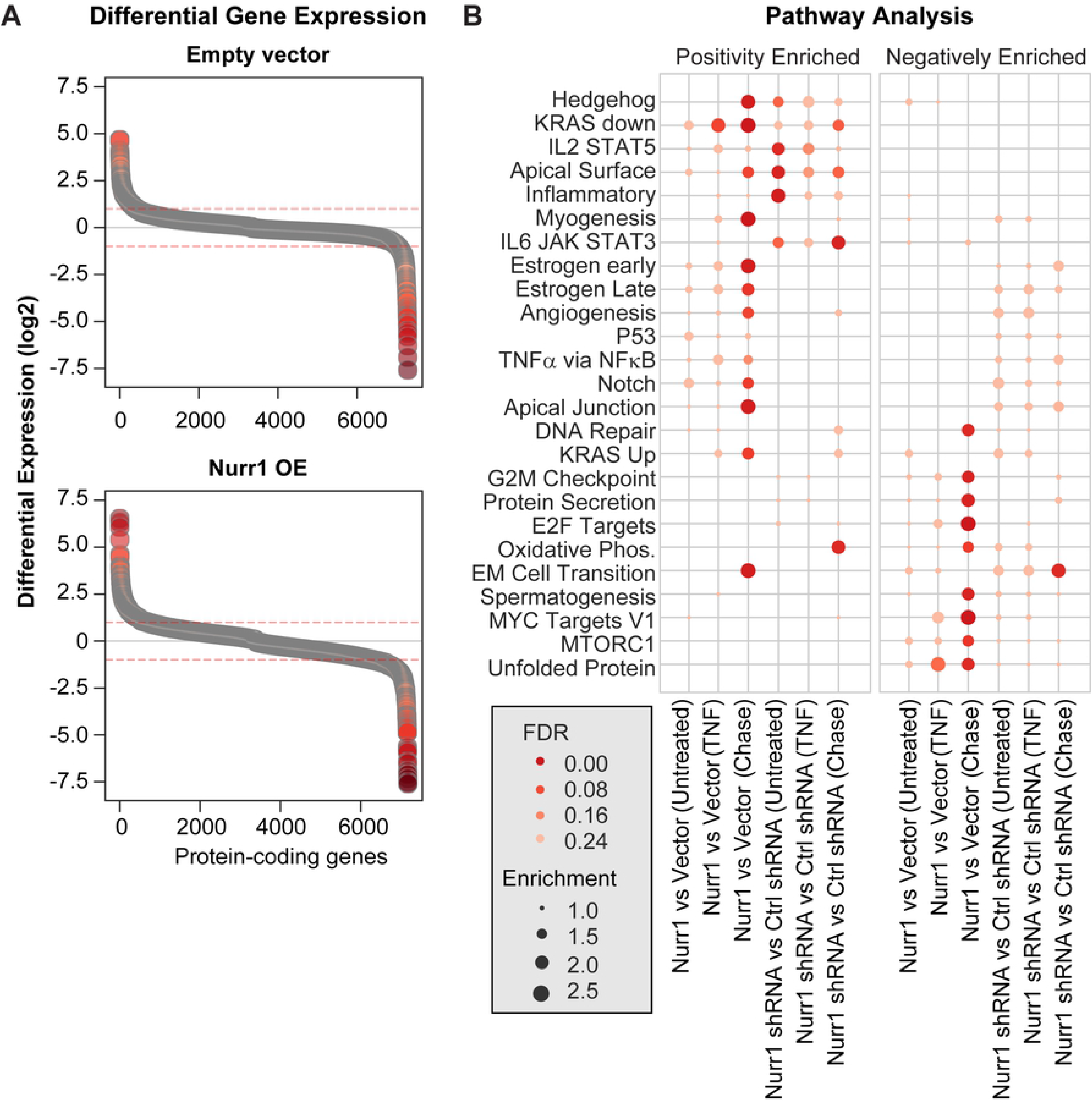
Nurr1 overexpression leads to the inhibition of critical cellular proliferation pathways. **A,** Patterns of differential gene expression during the chase step in vector-infected (top) and Nurr1 overexpressing (Nurr1 OE) cells. Dotted lines indicate the two-fold cut off level. **B,** Pathway analyses of Nurr1 overexpression at baseline, during TNF-α stimulation, and following the recovery period after TNF-α stimulation. The identities of specific highly enriched pathways are shown on the Y axis, and the comparisons are shown at the bottom. The color and size of circles correspond to statistical significance, as shown by FDR, and normalized enrichment values, respectively. Positive and negatively enriched pathways are shown in the left and right plot, respectively.

It is important to note that Nurr1 overexpression did not significantly interfere with the TNF-α signaling pathway during any step of these experiments (**Fig 5B**), suggesting that the cellular proliferation pathways we have identified are directly regulated by Nurr1. To further address this issue and determine whether Nurr1 simply accelerated the reversal of the normal microglial response to TNF-α stimulation during the chase, or if it regulated a distinct set of genes and pathways, we performed a gene trajectory analysis (**Fig 6A, S2 Fig**).

**Figure 6.**
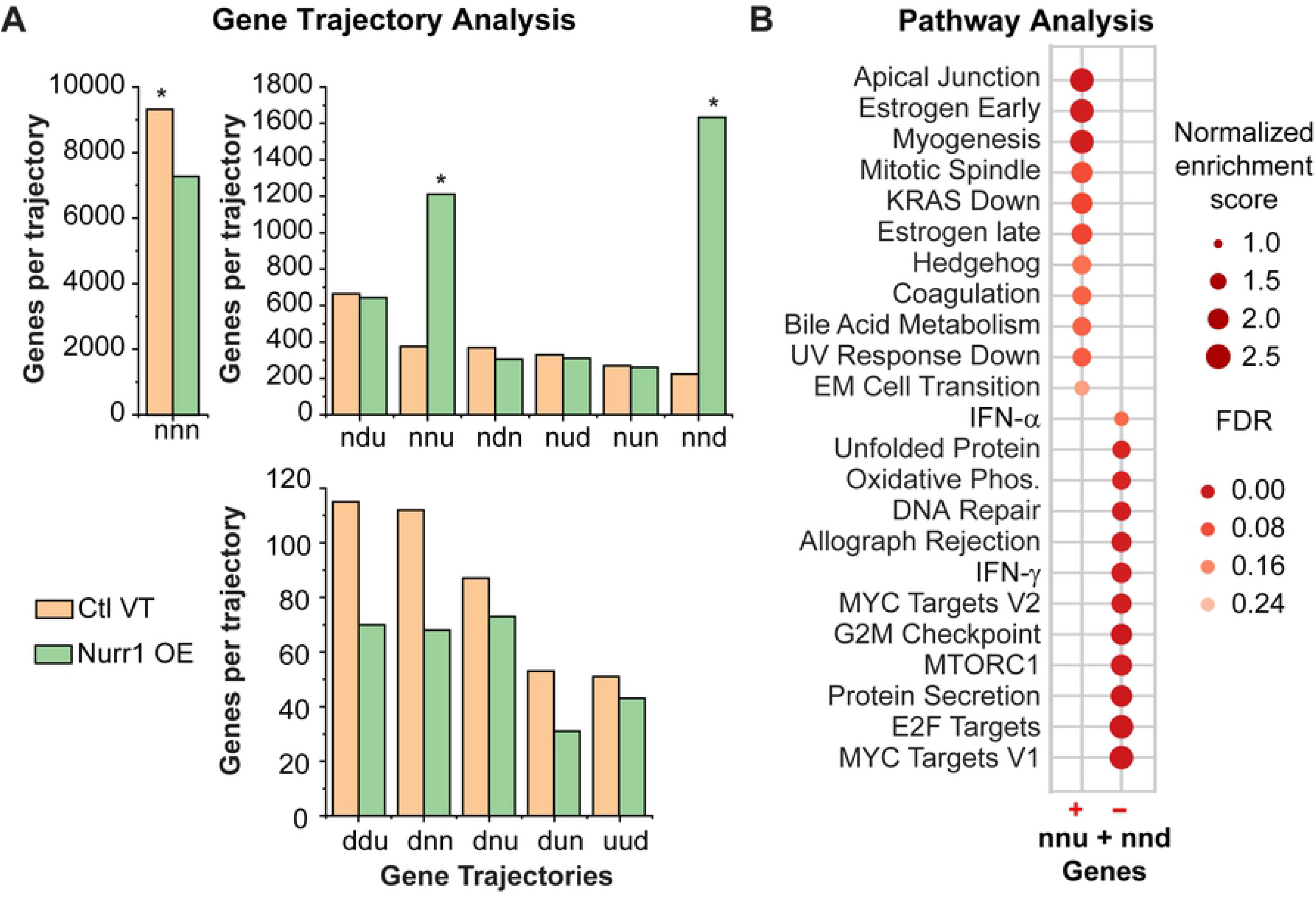
Nurr1 overexpression accelerates homeostasis of activated microglial cells by shutting down pathways involved in the maintenance of cellular activation and inflammation. **A,** Identification of genes selectively altered as a result of Nurr1 overexpression (Nurr1 OE), compared to the control empty vector (Ctl VT) cells, by trajectory analysis. Genes that are unaltered (n), downregulated (d) or upregulated (u) were identified during the activation and the chase steps and were clustered into families with similar profiles. The total number of genes in each category is indicated for both the control and Nurr1-overexpressing cells. Note that the major differences in the gene expression profiles are seen in genes that are either upregulated or downregulated during the chase (highlighted by asterisks). To enable the visualization of the trajectories with low, medium and high membership, the X axis for each group is shown separately. **B,** Pathway analysis using the Hallmark gene lists of the MSigDB database was performed on non-TNF-α-responsive genes that are exclusively altered in expression during the chase step in Nurr1 overexpressing cells, corresponding to genes which follow nnu and nnd trajectories in Nurr1 cells and an nnn trajectory in control cells (see **S3 Fig**). The identity of each pathway is shown to the left, and the direction of enrichment (+ or -) is shown at the bottom. The color and size of circles corresponded to statistical significance, as shown by FDR, and normalized enrichment values, respectively.

For the trajectory analysis we included RNA-Seq data from cells that were treated with the low dose of TNF-α (20 pg/ml), to simulate a sub-optimal activation signal. A pseudo-trajectory was defined containing three steps: Step 1 defines the changes in gene expression following stimulation with low dose TNF-α compared to untreated cells. Step 2 defines additional changes after stimulation with high dose TNF-α compared to cells treated with low dose TNF-α. Step 3 defines the gene expression changes following the chase step compared to cells treated with high dose TNF-α (**S2 Fig**). For each of these steps we calculated whether the expressed protein-coding genes were either upregulated (designated as “u”), downregulated (designated as “d”) or did not show differential expression in a statistically significant manner (designated as “n”). Genes that showed similar patterns of changes during each step were placed in the same category and named according to their pattern of change during these treatment steps. For example, those that did not show a change after low dose TNF-α stimulation (thus marked as n for Step 1), but were downregulated after high dose TNF-α treatment compared to cells treated with low dose TNF-α (marked as d for Step 2), and showed upregulation during the chase study compared to cells treated with high dose TNF-α (marked as u for step 3), were therefore designated as ndu.

Most genes did not show any change in their expression following the above treatments (designated as the “nnn” group) in both control (vector) and Nurr1-overexpressing cells (**Fig 6A**, **S2 Fig**), and as expected, control cells had higher numbers of nnn group genes than Nurr1 overexpressing cells.

Among those genes that showed an expression change in Nurr1 overexpressing cells, the majority belonged to genes that were not differentially expressed after either a low or high dose TNF-α treatment and exclusively changed their expression profiles during the chase step (i.e., nnu and nnd trajectories, **Fig 6A**, **S2 Fig**). We also noted that the number of genes in these two trajectories were markedly higher in Nurr1 overexpressing cells compared to control cells (i.e., over 800 and 1400 genes for nnu and nnd trajectories, respectively) while the number of genes in other trajectories with the exception of nnn differed by less than 100 genes (**Fig 6A**, **S2 Fig**).

We next confirmed that the genes showing the nnu and nnd trajectories in Nurr1 overexpressing cells were derived from a subset of the nnn trajectory genes in the control group and were therefore exclusively altered during the chase step in Nurr1 overexpressing cells. To further characterize the Nurr1-specific changes in expression patterns, we used the list of genes in each of the trajectories identified in Nurr1 overexpressing cells and defined their trajectory in control cells (**S3 Fig**). This analysis showed that over 1400 and ∼800 of the genes that fall into the nnd or nnu trajectories in Nurr1 overexpressing cells, respectively, have the nnn trajectory in control cells (**S3 Fig**). Thus, the main transcriptomic outcome of Nurr1 overexpression compared to control cells is the induction of changes in expression of a group of genes exclusively during the chase step. Importantly, this group of genes are not differentially expressed in the control cells during either of the three steps of these studies, nor during the TNF-α stimulation steps in Nurr1 overexpressing cells and therefore, the action of Nurr1 during the chase step does not correspond to a reversal of the TNF-α-induced changes.

In order to define the functional impact of this Nurr1-specific set of genes, we performed pathway analysis on the subset of genes that had either nnu or nnd trajectories in Nurr1 overexpressing cells, and a nnn trajectory in control cells (**Fig 6B**). Strikingly, these Nurr1-induced changes in gene expression during the chase step once again highlighted the downregulation of several key proliferative pathways, including: MYC, E2F and MTORC signaling, G2M checkpoint regulation, metabolic pathways such as oxidative phosphorylation, and inflammatory pathways such as IFN-α and IFN-γ response pathways (**Fig 6B**).

Heat maps of the differentially expressed genes further emphasized that the vast majority of genes in each pathway were downregulated in Nurr1 overexpressing cells. For example, among 69 and 60 represented MYC and E2F target genes, 66 and 54 were downregulated in Nurr1 overexpressing cells, respectively (**S4 Fig**). Finally, another compelling way of visualizing these results is to apply the pattern of expression of the Nurr1-specific genes to the KEGG cell cycle pathway (**S5 Fig**). The strong downregulation by Nurr1 at multiple steps in the cell cycle control pathway is immediately obvious.

Finally, we note as another measure of the specificity of the Nurr1 pathway, that the most enriched transcription factor binding motifs in proximity of the promoters of differentially expressed genes following TNF-α stimulation all display promoter motifs consistent with TNF-α activation (**S6 Fig)**.

Thus, the main impact of Nurr1 on the overall cellular response to inflammatory cytokines, in this case TNF-α, was to accelerate the cellular return to homeostasis by shutting down pathways involved in inflammation and microglial activation. While these anti-inflammatory, pro-homeostasis effects could indirectly lead to HIV proviral transcriptional shutdown, the enhanced downregulation of HIV expression in Nurr1 overexpressing cells, even under basal untreated conditions (**Fig 3D**), suggests that in addition to its pro-homeostasis effects, Nurr1 may also directly regulate the expression of the HIV provirus.

### Nurr1 promotes the recruitment of the CoREST/HDAC1/G9a/EZH2 repressor complex to the HIV promoter

Previous studies demonstrated that Nurr1 interacted with the corepressor 1 of REST (CoREST) repressor complex [50, 78]. The CoREST complex is comprised of multiple components including CoREST, repressor element-1 silencing transcription factor (REST), HDAC1/2, euchromatic histone lysine N-methyltransferase 2 (EHMT2), also known as *G9a*, lysine (K)-specific demethylase 1A (KDM1A), and enhancer of zeste 2 polycomb repressive complex 2 subunit (EZH2) [79, 80]. In microglial cells and astrocytes, after stimulation with lipopolysaccharide (LPS), Nurr1 promoted recruitment of this complex to the promoters of inflammatory genes such as IL-1β leading to epigenetic silencing. We postulated that the Nurr1/CoREST repression pathway might therefore also lead to direct regulation of HIV silencing as illustrated in **Fig 7A**.

**Figure 7.**
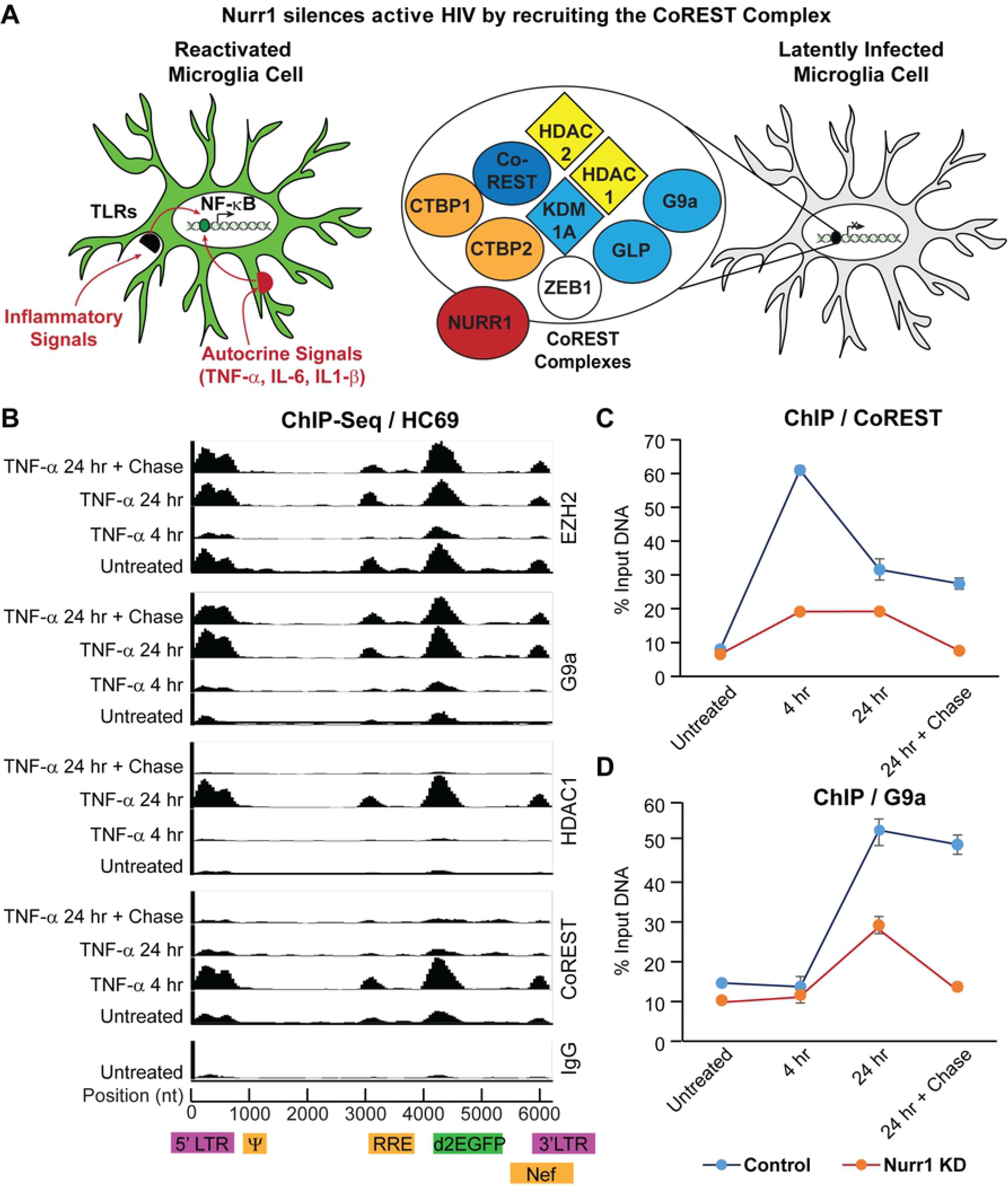
Nurr1 promotes recruitment of the CoREST repressor complex to HIV promoter. **A,** Schematic illustration of Nurr1-mediated epigenetic silencing of active HIV in microglial cells by recruiting the CoREST/HDAC1/G9a/EZH2 repression complex to HIV promoter. **B,** ChIP-seq signals (numbers of sequence reads on Y axis) along the reporter HIV-1 pro-viral genome (Figure 1A) on the X axis, resulting from ChIP-seq analysis with antibodies to EZH2, G9a, HDAC1, CoREST, and control IgG, respectively, and sheared chromatins prepared from HC69 cells that were un-treated, induced with TNF-α (400 pg/ml) for 4 hr and 24 hr respectively, or used in a chase experiment by continuously culturing HC69 cells in the absence of TNF-α for 24 hr after stimulating the cells with TNF-α (400 pg/ml) for 24 hr and washing with PBS. Construction of ChIP-seq DNA libraries with the ChIP products, enrichment for HIV-1 specific sequences, and data analysis following Ion Torrent sequencing were described in Materials & Methods. Positions of ChIP sequence reads along the viral genome were marked. **C & D**, levels of CoREST (C) and G9a (D) in HIV 5’LTR (+30 to +134) in HC69-control shRNA (Control) and HC69-Nurr1 shRNA (Nurr1 KD) cell lines that were treated as described in B. The levels of CoREST and G9a in HIV 5’LTR were measured by qPCR and calculated as percentages of the amounts of ChIP products over input DNA from triplicate qPCR.

To test this hypothesis, we first conducted co-immunoprecipitation (Co-IP) assays to confirm the association of Nurr1 with the CoREST repressor complex in HC69 cells (**S7 Fig.**). HC69-3X-FLAG-vector and HC69-3X-FLAG-Nurr1 cells were treated with and without a high dose of TNF-α for 4 hr (400 pg/ml) or 24 hr. After 24 hr TNF-α treatment the cells were chased in the absence of TNF-α for a further 24 hr. Total protein lysates from the differently treated cells were immunoprecipitated using a mouse monoclonal anti-FLAG antibody conjugated to magnetic beads. The anti-FLAG beads pulled down not only FLAG-tagged Nurr1 but also CoREST, HDAC1, G9a, and EZH2 from the HC69-3X-FLAG-Nurr1 cell lysates, demonstrating that in the microglial cells Nurr1 bound directly to the CoREST repressor complex. Notably, the amount of CoREST associated with Nurr1 increased after the cells were stimulated with TNF-α. In contrast, the amounts of G9a and EZH2 proteins associated with Nurr1 decreased at 4 hr post-TNF-α stimulation but rebounded at 24 hr post-TNF-α stimulation. Together, these results suggested that the Nurr1/CoREST/HDAC1/G9a/EZH2 complex were most likely dissociated from each other during early time points of TNF-α stimulation but were reassembled at later time points.

We next conducted ChIP-Seq experiments to demonstrate recruitment of the CoREST repressor complex to the activated HIV promoter in microglial cells. As shown in **Fig 7B,** CoREST, HDAC1, G9a, and EZH2 were all detected on the HIV provirus and were enriched near the promoter region following TNF-α activation. However, the recruitment kinetics of each component was distinct, with CoREST being recruited to HIV promoter during early time points of TNF-α exposure and HDAC1, G9a, and EZH2 being recruited at late time points. These results are consistent with the Co-IP results shown in **S7 Fig**. Specifically, the levels of CoREST at the HIV promoter peaked at 4 hr post-TNF-α stimulation and decreased at 24 hr post-treatment, whereas the levels of G9a, EZH2, and HDAC1 at the HIV promoter decreased at 4 hr post-TNF-α stimulation when compared to un-treated cells. However, these epigenetic silencers returned to HIV promoter in a much more robust manner at 24 hr post-stimulation.

To provide direct evidence that Nurr1 mediates the recruitment of the CoREST/HDAC1/G9a/EZH2 complex to HIV promoter, we treated HC69 cells expressing control shRNA and Nurr1 shRNA with high dose TNF-α, followed by a 24 hr chase. We then conducted additional ChIP experiments and measured the ChIP products by quantitative PCR (qPCR). As shown in **Fig 7C**, CoREST was strongly recruited to HIV promoter at 4 hr post TNF-α stimulation in HC69 cells expressing control shRNA, however, its recruitment was substantially inhibited in Nurr1 KD cells. Similarly, G9a level in HIV promoter peaked at 24 hr post TNF-α stimulation in HC69 cells expressing control shRNA but its recruitment was also reduced in Nurr1 KD cells. Taken together, these results clearly demonstrated a pivotal role for Nurr1 in mediating recruitment of the CoREST/HDAC1/G9a/EZH2 repressor complex to the promoter of active HIV for epigenetic silencing consistent with the model shown in **Fig 7A**.

### The CoREST/HDAC1/G9a/EZH2 repressor complex silences HIV in microglial cells

To further investigate how the CoREST/HDAC1/G9a/EZH2 complex contributes to HIV silencing, we treated HC69 cells with high dose (400 pg/ml) TNF-α for 24 hr followed by a chase in the absence or presence of epigenetic inhibitors that target the CoREST complex, specifically: HDAC inhibitor suberoylanilide hydroxamic acid (SAHA), G9a inhibitor UNC0638, and EZH2 inhibitor GSK343. The numbers of GFP+ cells dropped from 88.4% to 67.03% at 48 hr after TNF-α withdrawal when cells were cultured in the absence of the inhibitors (**Fig 8A**). However, in the presence of SAHA, UNC0638, or GSK343, the numbers of GFP+ cells remained higher (i.e., 77.8%, 85.5%, and 84.7% respectively), indicating that functional inhibition of these epigenetic silencers prevented active HIV from reverting to latency.

**Figure 8.**
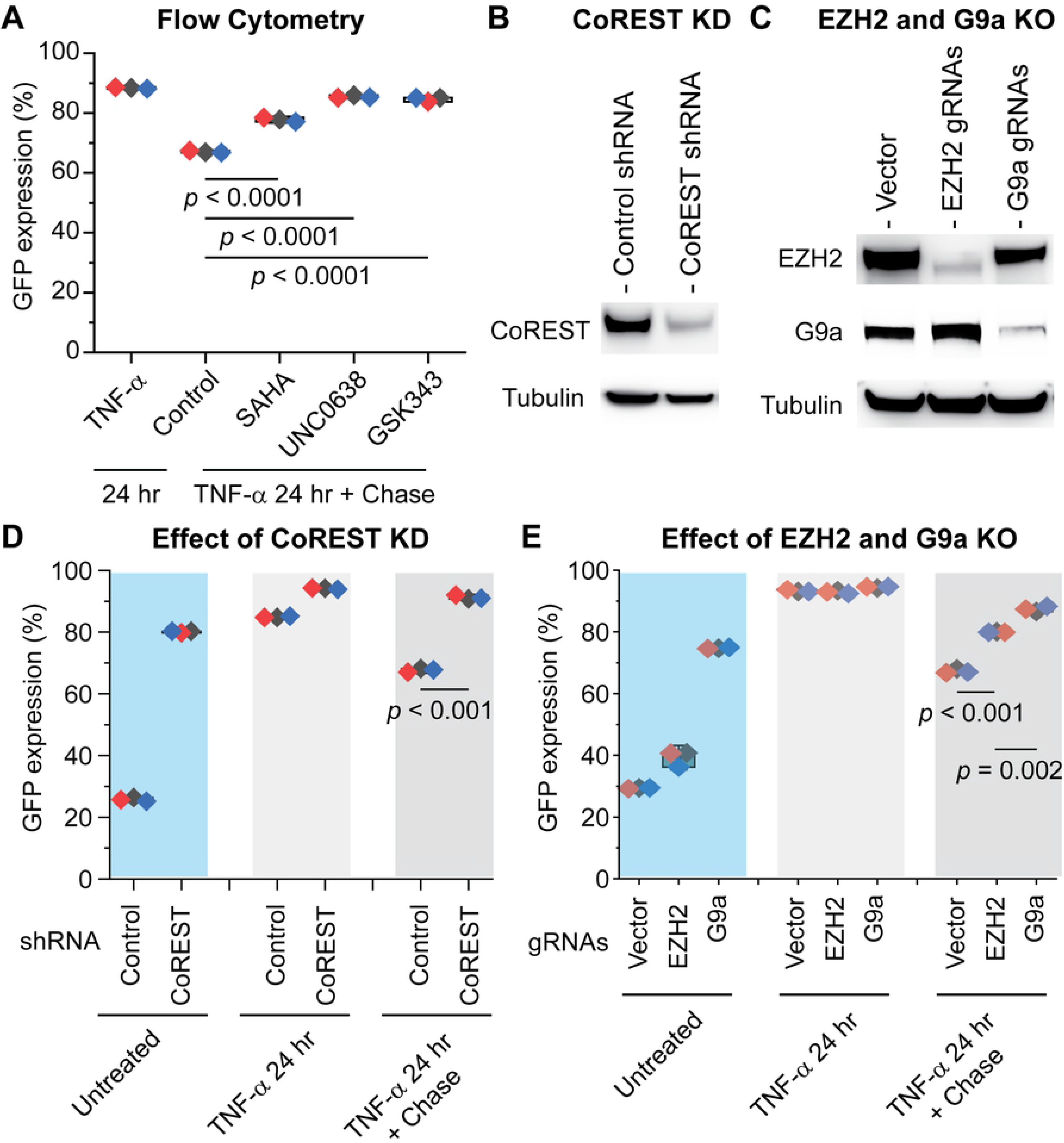
The CoREST repressor complex plays a pivotal role in silencing active HIV in microglial cells. **A,** Inhibition of HDAC1, G9a, and EZH2 blocked silencing of activated HIV in HC69 cells. HC69 cells were stimulated with high dose (400 pg/ml) TNF-α for 24 hr. After washing with PBS, the cells were cultured in the presence of DMSO (placebo, Control), HDAC inhibitor SAHA (2 μM), G9a inhibitor UNC0638 (2.5 μM), and EZH2 inhibitor GSK343 (2.5 μM), respectively, for 48 hr. The levels of GFP expression for each treatment were measured by flow cytometry and calculated from three independent experiments, with *p* values between the control and treatment with each inhibitor indicated. **B,** Verification of CoREST KD by Western blot detection of CoREST protein expression in HC69 cell lines stably expressing control shRNA or CoREST-specific shRNA. **C,** Verification of EZH2 and G9a KO by Western blot detection of G9a and EZH2 protein expression in HC69 cells stably expressing CRISPR/Cas9 and G9a or EZH2 specific gRNA, which were compared to the control HC69 cells stably expressing CRISPR/Cas9 without gRNA. β-tubulin was used as a loading control for all Western blot analysis. **D,** CoREST KD prevents HIV silencing. The HC69-control shRNA and HC69-CoREST-shRNA cells were untreated, induced with high dose (400 pg/ml) TNF-α for 24 hr, or used in a chase experiment by continuous culturing the cells for 48 hr after TNF-α stimulation for 24 hr and washes with PBS. GFP expression levels of all cells were measured by flow cytometry and the mean values were calculated from three independent experiments. Significant differences were observed between the HC69-control shRNA and HC69-CoREST shRNA cell lines. **E,** G9a and EZH2 KO prevents HIV silencing. Evaluation of the HC69 cell lines expressing G9a or EZH2 specific gRNA or empty vector by flow cytometry following the same protocol as in panel D. There was a significant difference between HC69-vector and HC69 EZH2 or G9a KO cell lines at 48 hr after TNF-α withdrawal, with *p* < 0.01.

To confirm the role of these epigenetic silencers, we generated HC69 cell lines stably expressing CoREST-specific shRNA or CRISPR/Cas9/guide RNA (gRNA) for G9a or EZH2. We confirmed successful KD or knock out (KO) of these proteins in these cell lines by Western blot analysis (**Fig 8B** & **C**). The genetically modified cells were activated with a high dose of TNF-α (400 pg/ml) for 24 hr, followed by culturing the cells in the absence of TNF-α for 48 hr and measurement of GFP expression. CoREST KD substantially increased GFP expression (80.1% GFP+ vs. 25.8% in control cell) even without TNF-α stimulation (**Fig 8D**). Stimulation with high-dose TNF-α for 24 hr resulted in 94.1% and 84.9% GFP+ cells in CoREST KD and control cells respectively. However, after TNF-α withdrawal and subsequent culture for 48 h, the numbers of GFP^+^ cells decreased significantly in cells expressing control shRNA (67.7%) but remained high in CoREST KD cells (91.3%), confirming that CoREST was crucial for the silencing of active HIV in microglial cells. Similar results were also seen with the G9a and EZH2 KO cell lines (**Fig 8E**).

Therefore, both the ChIP experiments and gene knockout results demonstrate a pivotal role for the CoREST/HDAC1/G9a/EZH2 transcription repressor complex in silencing active HIV in microglial cells.

### Nurr1 regulates HIV in iPSC-derived microglial cells

Finally, to confirm that Nurr1 is also critical for the silencing of HIV in primary microglial cells, we infected iPSC-derived human microglial cells (iMG) with the same HIV reporter virus described earlier (**Fig 1A)**. About 50% of the iMG became GFP+ two days after HIV infection (**Fig 9A**). We then treated the infected iMG with 6-MP and another Nurr1 agonist, amodiaquine (AQ) [56, 67], for four days. Both 6-MP and AQ decreased the number of GFP+ cells in a dose-dependent manner (**Fig 9B & C**) and lowered the levels of HIV un-spliced transcripts (**Fig 9D**). Both agonists also dose-dependently reduced MMP2 mRNA in iMG (**Fig 9E**). Collectively, results from both hµglia and iMG strongly suggested an important role for Nurr1 in HIV silencing in microglial cells.

**Figure 9.**
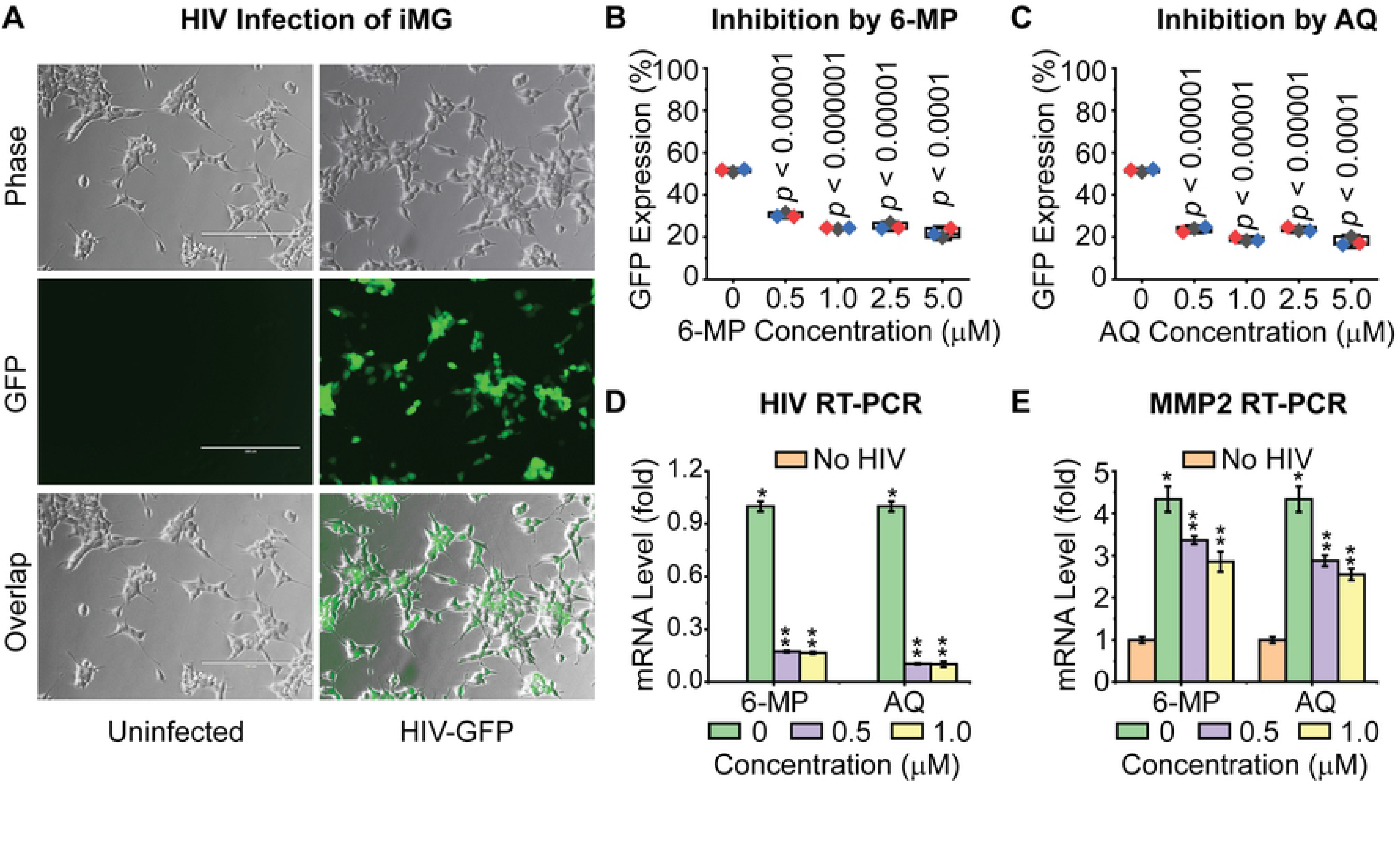
Nurr1 Mediates HIV silencing in iPSC-derived microglial cells (iMG). **A,** Representative phase contrast, GFP, and overlapped images of iMG that were un-infected or infected with the reporter HIV-1 shown in Fig 1A, at 48 hr post-infection (hpi). HIV-infected iMG were treated with different doses of Nurr1 agonist 6-MP or AQ for four days, followed by flow cytometry analysis of GFP expression. **B,** The average levels of GFP expression in iMG treated with various doses of 6-MP. **C,** The average levels of GFP expression in iMG treated with various doses of AQ were calculated from three replicates. **D,** The levels of HIV RNA (un-spliced) in the cells described in panels A and B, were measured by RT-qPCR. **E,** The mRNA level of Nurr1 target gene MMP2 in the same cells was measured by qRT-PCR. The average levels of HIV transcript and MMP2 mRNA in each sample were calculated from triplicates of qRT-qPCR. Differences in HIV and MMP2 mRNA levels between un-treated cells and cells treated with different doses of 6-MP or AQ were statistically significant (** *p*-values <0.001). HIV transcripts were only detected in infected iMG cells (panel D). MMP2 mRNA was significantly elevated in HIV infected iMG (panel E).

## DISCUSSION

### Epigenetic control of HIV latency in microglial cells

Microglial cells are one of the major cellular reservoirs of HIV in the central nervous system (CNS) [4, 5]. These long-lived cells contribute to increased neuroinflammation and oxidative stress [4, 81], and development of HAND by secreting a variety of neurotoxins as well as harmful HIV proteins such as gP120, Tat, Rev, etc. [82, 83]. Eradication or complete silencing of HIV-infected microglial cells is therefore crucial not only for an HIV cure, but also to prevent the development of HAND, which affects the majority of HIV infected individuals.

Previous studies involving HIV-1 infection of transformed cell lines suggested that epigenetic regulation plays a major role in the establishment and persistence of HIV latency in astrocytes and microglial cells [84, 85]. The cellular COUP transcription factor (COUP-TF) interacting protein (CTIP2) forms a large transcriptional repressor complex with epigenetic silences including the histone deacetylases HDAC1/2, the histone methyltransferases SUV39H1 and SET1, the lysine(K)-specific demethylase KDM1, and heterochromatin protein1 (HP1) [86, 87]. Recruitment of this complex to HIV-1 promoter leads to proviral genome silencing due to reduced histone acetylation and increased levels of histone 3 tri-methylations at lysine 9 (H3K9me3) [86–89]. At the same time, CTIP2 forms another complex with CDK9, Cyclin T1, HEXIM1, 7SK snRNA, and high mobility group AT-hook 1 (HMGA1), which is also recruited to HIV-1 promoter [87, 89]. In the absence of HIV-1 Tat, this complex with inactive pTEFb further supports HIV-1 latency by preventing elongation of RNA polymerase II for active transcription [90]. Nevertheless, it remains unknown if these mechanisms also apply to HIV-infected primary microglial cells as transformed cells often behave quite differently.

In a previous study [37], we demonstrated that autocrine inflammatory cytokines such as TNF-α were major drivers for spontaneous HIV reactivation in microglial cells, and activation of GR with its specific ligands such as dexamethasone antagonized the effects of cytokines on HIV reactivation [45]. However, we observed that the reactivated HIV was subsequently silenced in microglial cells in the absence of dexamethasone, suggesting the existence of additional HIV silencing mechanisms.

### Silencing of HIV by Nurr1 and CoREST

In the present study, we identified the nuclear receptor Nurr1 as a key HIV silencing factor. Overexpression of Nurr1 had little effect on preventing reactivation of latent HIV but strongly enhanced silencing of active HIV after TNF-α stimulation and subsequent withdrawal. Inversely, KD of endogenous Nurr1 in HC69 cells inhibited silencing of active HIV after TNF-α withdrawal. Thus, results from both overexpression and KD experiments unequivocally demonstrated a pivotal role of Nurr1 in silencing active HIV. Mechanistically, we demonstrated that Nurr1 interacted with the CoREST/HDAC1/G9a/EZH2 repressor complex as reported previously for cellular early response genes [50]. Nurr1 promoted the recruitment of CoREST complexes to the HIV promoter following TNF-α stimulation and subsequent withdrawal.

These epigenetic silencers likely silence active HIV by promoting histone de-acetylation and repressive di- or tri-methylations. Consistent with this hypothesis, functional inhibition with specific inhibitors or expressional KD or KO of each component of the repressor complex including CoREST, G9a, and EZH2 strongly inhibited HIV silencing. Data from RNA-Seq analysis indicated that Nurr1 might also utilize this “post-TNF-α stimulation” epigenetic silencing mechanism to repress many host genes.

### Regulation of HIV latency in microglial cells by nuclear receptors

Nuclear receptors are special transcription factors that turn on or turn off expression of target genes upon specific ligand binding [91]. Accumulating evidence suggest that nuclear receptors play an important role in regulating HIV expression. For example, estrogen receptor (ER) and GR have been found to promote HIV latency in T cells and microglial cells respectively [45, 63]. In this study, by using both immortalized and iPSC-derived human microglial cells, we provided comprehensive data to demonstrate that Nurr1 promoted HIV latency by silencing active HIV. In contrast, we did not see significant effects of Nur77 and Nor1 overexpression on HIV when HC69 cells were stimulated with TNF-α. However, we have not examined possible effects of these nuclear receptors on HIV when microglial cells are activated through other signaling pathways such as the “Toll-like” receptor signaling pathway [38]. In addition, it is well known that the different Nerve Growth Factor IB-like nuclear receptors interact with each other or with other nuclear receptors such as GR and the retinoid X receptors (RXR) [92, 93]. Therefore, in future experiments, we plan to investigate how Nur77, Nurr1, and Nor1 impact HIV expression in microglial cells in response to different stimuli and whether they exert any synergistic effect on HIV expression between themselves or with other interacting nuclear receptors.

### Role of Nurr1 in maintaining brain homeostasis

The roles of Nerve Growth Factor IB-like nuclear receptors in brain development and homeostasis are well established. Both Nor1 and Nurr1 are essential for differentiation and survival of dopaminergic neurons [46–49]. Nurr1 deficiency in embryonic ventral midbrain cells results in their failure to migrate and innervation of their striatal target areas [94, 95]. Nurr1 deficiency or reduced expression due to mutations in adults is a major contributing factor in the pathogenesis of Parkinson’s disease [96]. Nurr1 is also expressed in non-neuronal cells including monocytes, macrophages, microglia, and astrocytes. Its expression is reduced in the peripheral blood lymphocytes (PBL) of patients with Parkinson’s disease compared with healthy controls [97]. Lower levels of Nurr1 in the brain and blood represents increased risks of Parkinson’s disease and other neurodegenerative diseases in adults [97].

Nurr1 protects dopaminergic neurons from inflammation-induced neurotoxicity through the inhibition of pro-inflammatory mediator expression in microglia and astrocytes by recruiting CoREST corepressor complexes to NF-κB target genes [50, 98]. A reduction of Nurr1 expression in neurons does not affect their death but enhances expression of inflammatory mediators, and the survival rate of neurons decreases in response to inflammatory stimuli in the Nurr1 deficiency condition [50]. Multiple studies reported that activation of Nurr1 reduces inflammation, protects neurons, and decreases Parkinson’s disease related symptoms [53, 65, 67, 99].

Although the pathogenesis of Parkinson’s disease and other types of neurodegeneration remains obscure, increasing evidence suggests that inflammatory responses are responsible for the progression of most neurodegenerative diseases [100]. These responses include accumulation of inflammatory mediators, such as inflammatory cytokines and proteases in the substantia nigra and the striatum, as well as activation of the microglia [101], which are also common features of HAND [4, 102].

### Anti-inflammatory role for Nurr1 in HIV-infected microglial cells

Little is known on how HIV infection impacts expression or functionality of Nurr1 and other nuclear receptors in the brain. Microglial activation is triggered by a series of neurochemical mediators such as IFN-γ, inducible nitric oxide synthase (iNOS), IL-1β, and TNF-α [103–106]. HIV infection of the brain likely further increases the levels of these mediators. Interestingly, data from our RNA-Seq experiments reveal that Nurr1 overexpression pushed the activated microglial cells towards homeostasis following TNF-α stimulation and subsequent withdrawal by repressing NF-κB signaling pathway and genes involved in cellular activity and IFN-α and INF-γ responses. Thus, in addition to silencing HIV, Nurr1 apparently plays a crucial role in suppression of microglia activation. This finding is consistent with a recent report that glycolysis downregulation is a hallmark of HIV-1 latency in microglial cells [107].

Further studies are warranted to determine the expression levels of these nuclear receptors in HIV patients and investigate whether their deficiency or malfunction contributes to development of HAND. Interestingly, multiple Nurr1 agonists exhibit strong therapeutic effects and potentials for Parkinson’s disease in pre-clinical animal study and human trials [108]. In this study, we tested the Nurr1 agonists 6-MP and AQ. Both agents strongly inhibited expression of HIV and the neurotoxin MMP2 in HC69 cells and iMGs. In future studies, it would be of great interest to test additional agonists, particularly those new generations of Nurr1 agonists currently on pre-clinical and human trials, for their anti-HIV activity and eventual application in the clinic for treatment of HAND.

## MATERIALS & METHODS

### Chemicals and reagents

TNF-α (Invitrogen, Cat. #PHC3015) was used to induce HIV-1 reactivation in microglial cells. Nor1 and Nurr1 agonists 6-mercaptopurine (6-MP) (Millipore-Sigma, Cat#38171) and amodiaquine (AQ) (Millipore-Sigma, Cat#SMB00947) were used to activate the Nerve Growth Factor IB-like nuclear receptors. GSK343 (Sigma Aldrich, Cat# SML0766), UNC0638 (Sigma Aldrich, Cat#U4885), and suberoylanilide hydroxamic acid (SAHA, Millipore-Sigma, Cat#SML0061) were used to examine the effects of EZH2, H9a, and HDAC1/2 on HIV silencing respectively.

Numerous antibodies were used for Western blot analysis, co-immunoprecipitation (Co-IP), and chromatin immunoprecipitation (ChIP) assays, including a mouse monoclonal anti-FLAG M2 antibody (Sigma, Cat# F1804), a rabbit polyclonal anti-Nurr1 antibody (Sata Cruz Biotechnology, Cat# sc-991), a mouse monoclonal anti-Nurr1 antibody (Santa Cruz Biotechnology, Cat# sc-81345), a mouse monoclonal anti-Nor1 antibody (Perseus Proteomics, Cat# PP-H7833-00), a rabbit polyclonal anti-Nur77 antibody (Cell Signaling, Cat# 3960S), a rabbit monoclonal anti-MMP2 antibody (Cell Signaling, Cat#40994), a rabbit polyclonal anti-CoREST antibody (EMD Millipore, Cat# 07-579), a rabbit polyclonal anti-HDAC1 antibody (Santa Cruz Biotechnology, Cat# sc-7872), a rabbit polyclonal anti-G9a antibody (Cell Signaling, cat#3306S), a rabbit polyclonal anti-EZH2 antibody (Cell Signaling, Cat#5246S), a rabbit polyclonal anti-acetylated histone 3 (Ac-H3) antibody (Cell Signaling, Cat#9677S), a rabbit polyclonal anti-H3K27me3 antibody (EDM Millipore, Cat#07-449), a rabbit monoclonal anti-H3K27me2 antibody (Cell Signaling, Cat#9728S), a mouse monoclonal anti-HIV Nef antibody (Abcam, Cat#ab42355), and a mouse monoclonal anti-RNA polymerase II antibody (Abcam, Cat#ab817).

### Cells and flow cytometry analysis of HIV/GFP expression

HIV-1 infected immortalized human microglial (hµglia) HC69 cells were cultured and maintained as described as previously [37]. Induced pluripotent stem cells (iPSC)-derived human microglial cells (iMG) (Tempo Bioscience, Cat#SKU 1001.1) were plated, allowed to differentiate and maintained in culture on plates pre-coated with Matrigel matrix (Corning, Cat#356254) according to the manufacturer’s instructions. The iMG were infected with EFGP HIV-1 reporter virus at 1 to 1 (cell-to-virus moiety), which was produced, purified, and titrated as described previously [37]. Two days after infection, the iMG were treated with and without the Nurr1 agonists 6-MP and AQ for four days. Infected with the same EGFP-reporter HIV-1 virus (**Fig 1 A**), HIV expression in hµglia and iMG cells was measured and quantified with percentage (%) of GFP+ cells by flow cytometry as described previously [45].

### Lenti-viral construction and production, and generation of stable cell lines

Three lentiviral constructs, pLV[Bxp]-Bsd-CMV>3xFLAG-Nur77, pLV[Bxp]-Bsd-CMV>3xFLAG-Nurr1, and pLV[Bxp]-Bsd-CMV>3xFLAG-Nor1 were generated by inserting the full-length open reading frame (ORF) of human NR4A1 (Nur77), NR4A2 (Nurr1), and NR4A3 (Nor1) cDNA fragment into the empty vector pLV[Bxp]-Bsd-CMV>3xFLA immediately downstream of the Kozak sequence (VectorBuilder, vector ID: VB180227-1135bmn, VB180227-1134jht, and VB180227-1136rwc). The inserted cDNA was also “in frame” fused with the coding sequence of the N-terminal 3X-FLAG peptide tag, allowing to generate N-terminal 3xFLAG-tagged proteins. Two lentiviral constructs expressing human Nurr1-specific shRNA (5’GGTTCGCACAGACAGTTTAAA3’ and 5’ATACGTGTGTTTAGCAAATAA3’), one lentiviral construct expressing human CoREST-specific shRNA (5’CCCAATAATGGCCAGAATAAA3’), and two lentiviral constructs expressing control shRNAs (5’CCTAAGGTTAAGTCGCCCTCG3’ and 5’ CAACAAGATGAAGAGCACCA3’) were purchased from VectorBuilder. All lentiviral constructs carried an ampicillin resistance gene for selection in bacteria (*E. coli*) and a blasticidin resistance gene for selection of stable expression in mammalian cells. Infectious viral particles with each of these lentiviral constructs were produced by co-transfecting 293T cells with packaging plasmid psPAX2 (Addgene, Cat#12260) and Env Vector pCMV-VSVg (Addgene, Cat#138479). HC69 cells stably expressing 3X-FLAG-Nur77, 3X-FLAG-Nurr1, 3X-FLAG-Nor1, empty vector, gene-specific shRNA and control shRNA were generated by infection of the cells with purified lentiviral particles for two days, followed by culturing the cells in the presence of blasticidin at 10 μg/ml.

To investigate the effects of G9a and EZH2 on HIV silencing, we conducted CRISPR/Cas9 mediated “knocking out” (KO) of these genes in HC69 cells, using a dual CRISPR/Cas9 gRNA lentiviral vector. Two different guide RNAs targeting EZH2 (TGAGCTCATTGCGCGGGACT and GATCTGGAGGATCACCGAGA) or G9a (TTCCCCATGCCCTCGCATCC and GTGGCAGCCCCACGGCTGAA) were cloned into lentiCRISPR v2-Blast plasmid following the protocol described previously [109]. LentiCRISPR v2-Blast was a gift from Mohan Babu (Addgene plasmid # 83480). VSV-G pseudotyped viruses expressing CRISPR/Cas9 gRNAs were produced in HEK 293T cells by transfection of lentiCRISPR v2 plasmids together with psPAX2 and pCMV-VSV-G. HC69 cells infected with the EZH2 or G9a KO lentiviruses were cultured in the presence of blasticidin (10 μg/ml). Successful KO of these genes in HC69 cells were verified by Western blot analysis of EZH2 and G9a proteins in the resulting cell lines.

### Reverse transcription and quantitative polymerase chain reaction (RT-qPCR)

Total RNAs from HC69 or HIV-infected IMG cells with different treatments were isolated by using the RNeasy Plus Mini kit from Qiagen (Cat#74134). The purified total RNAs were converted to first-strand cDNAs by using a reverse transcription kit (Bio-Rad, Cat#1708891). The relative levels of HIV-1 un-spliced transcript and human MMP-2 mRNA were measured by qRT-PCR using the primers 5’AGGGACCTGAAAGCGAAAG3’ (HIV-1 un-spliced-forward) and 5’AATGATACGGCGACGACCNNNNNNNNNN3’ (HIV-1 un-spliced-reverse), and 5’ATAACCTGGATGCCGTCGT-3′ (MMP2 forward) and AGGCACCCTTGAAGAAGTAGC-3′ (MMP2 reverse), respectively. The mRNA level of the housekeeping gene β-actin in each sample was used as reference for normalization, TCCTCTCCCAAGTCCACACAGG-3′ (forward) and 5’-GGGCACGAAGGCTCATCATTC-3′ (reverse). Each qRT-PCR was conducted in triplicates.

### ChIP and ChIP-seq analyses

Standard procedures were followed for all ChIP assays. Briefly, cells were fixed with 1% Formaldehyde for 10 minutes (min) at room temperature, followed by incubation in PBS containing 125 mM glysine for 10 min at room temperature. After two washes with ice-cold PBS, cells were re-suspended and allowed to swell in CE buffer [10 mM Hepes, pH7.9, 60 mM KCl, 1 mM EDTA, 0.5% NP-40, 1 mM DTT] on ice for 10 min. After centrifugation at 2,000 g for 10 min at 4°C, nuclei were re-suspended in SDS lysis buffer [50 mM Tris-HCl, 1 mM EDTA, 0.5% SDS] and incubated on ice for 10 min. Sheared chromatins were prepared by sonicating the nuclei lysate to generate DNA fragments in the range of 250 to 500 bps. ChIP assays with specific antibodies were carried out in ChIP dilution buffer [16.7 mM Tris-HCl, pH 8.1, 167 mM NaCl, 1.2 mM EDTA, 1.1% Triton X-100, and 0.01% SDS] containing 5 μg antibody and 50 ul protein-A/protein-G magnetic beads per reaction at 4°C for overnight with rotation, followed by consecutive washes with low salt wash buffer [20mM Tris-HCl, pH8.1, 150 mM NaCl, 1 mM EDTA, 1% Triton X-100, 0.1% SDS], high salt wash buffer [20mM Tris-HCl, pH8.1, 500 mM NaCl, 1 mM EDTA, 1% Triton X-100, 0.1% SDS], and RIPA buffer [20 mM Tris-HCl, pH7.5, 150 mM NaCl, 5 mM EDTA, 0.5% Triton X-100, 0.5% sodium deoxycholate, and 0.1% SDS]. The washed beads were then re-suspended in elution buffer [50 mM Tris-HCl, pH 6.5, 20 mM NaCl, 100 mM NaHCO3, 1 mM EDTA, 1% SDS, 100 μg/ml proteinase K] and incubated at 50°C for 2h. Supernatants from the beads were collected and used for ChIP DNA purification using Qiagen’s PCR purification kit (Cat#28104). Quantification of input and ChIP DNA corresponding to HIV-1 promoter region was conducted by qPCR using specific primers as reported previously [110].

For ChIP-seq analyses, the DNA products from each ChIP assay were first end repaired with end repair enzyme mix (New England Biolabs, Inc., Cat#M6630), then ligated to NEBNext adaptor included in the NEBNext^®^ Ultra^™^ II DNA Library Prep Kit for Illumina^®^ (Cat#E7645L) according to the manufacturer’s instruction, followed by PCR amplification with a specific pair of bar-coded primers. Next, to enrich HIV-1 specific sequences in the library, DNA samples from all ChIP assays were pooled, denatured at 98°C for 10 min, and then subjected to hybridization with 50 times excessive amount of biotin-labelled and pre-denatured HIV-1 genomic DNA in hybridization buffer containing 5XSSC and salmon sperm DNA (100 μg/ml) at 65°C for 1 h. Fragments hybridizing to biotin-labelled HIV-1 DNA were pulled down by incubating the hybridization reaction with streptavidin-conjugated magnetic beads (ThermoFisher Scientific, Cat#88816) at room temperature for 30 min, followed by three times washes with ion wash buffer and elution in water. The enriched ChIP library DNA was PCR amplified with Ion A and Ion P1 primers, and PCR fragments in the range from 300 to 500 bps were purified from agarose gel after electrophoresis and loaded for Ion Torrent sequencing.

We aligned the sequence reads to NL4.3-Cd8a-EGFP-Nef+ HIV-1 genome. Raw fastq sequencing data were imported to *the public server at usegalaxy.org for analysis* [111]. We used FASTX-Toolkit for deconvolution of reads. Read mapping was performed by Bowtie2 tool with default settings using the NL4.3-Cd8a-EGFP-Nef+ HIV-1 as a reference genome [112, 113]. DeepTool2 was used to make graphs for distribution of mapped reads along HIV-1 genome [114].

### RNA-Seq and data analysis

Approximately 2 million GFP-negative cells from each of the cell lines HC69-3X-FLAG-vector, HC69-3X-FLAG-Nor1, HC69-3X-FLAG-Nurr1, HC69-control shRNA, and HC69-Nurr1 shRNA were collected from sorting. The isolated cells were expanded in DMEM culture media with low glucose (1g/L) and 1% FBS for 48 hr in the presence of dexamethasone (1μg/ml) to maintain HIV latency as reported previously [45]. The cells were next cultured in fresh medium without dexamethasone, un-treated, or treated with low dose (20 pg/ml) and high dose (400 pg/ml) TNF-α for 24 h. One portion of the cells treated with high dose TNF-α were washed twice with PBS, followed by culturing in fresh medium in the absence of TNF-α and dexamethasone for 48 h. Total RNAs from each cell line with different treatments were isolated by using the RNeasy Plus Mini kit from Qiagen (Cat#74134). The isolated RNAs were treated with RNase-free DNAse I at 37 °C for 30 min to remove genomic DNA, followed by a second-round purification using the same RNA purification kit. For reproducibility concerns, the RNA-Seq analysis consisted of RNA samples from two independent experiments performed several months apart.

Total cellular RNA was subjected to 150 base long, paired end RNA-Seq on an NovaSeq 6000 instrument. RNA-Seq reads were quality controlled using Fastqc and trimmed for any leftover adaptor-derived sequences, and sequences with Phred score less than 30 with Trim Galore, which is a wrapper based on Cutadapt and FastQC. Any reads shorter than 40 nucleotides after the trimming was not used in alignment. The pre-processed reads were aligned to the human genome (hg38/GRCh38) with the Gencode release 28 as the reference annotations using STAR version 2.7.2b [115], followed by gene-level quantitation using htseq-count [116]. In parallel, the pre-processed reads were pseudoaligned using Kallisto version 0.43.1 [117], with 100 rounds of bootstrapping to the Gencode release 28 of the human transcriptome to which the sequence of the transfected HIV genome and the deduced HIV spliced transcripts were added. The resulting quantitations were normalized using Sleuth. The two pipelines yielded concordant results. Pairwise differential expression tests were performed using generalized linear models as implemented in edgeR (QL) [118], and false discovery rate (FDR) values were calculated for each differential expression value.

Protein-coding genes that were expressed at a minimum abundance of 5 transcripts per million (TPM) were used for pathway analysis with fold change values as the ranking parameter while controlling false discovery rate at 0.05. Gene Set Enrichment Analysis (GSEA) package was used to identify the enriched pathway and promoter elements using mSigDB and KEGG databases. Pathways that showed an FDR q-value <= 0.25 were considered significantly enriched, per the GSEA package guidelines. The number of genes contributing to the enrichment score was calculated using the leading edge output of GSEA (tag multiplied by size).

### Identification of marker genes for each study group

After filtration of the raw reads to remove low quality reads and mapping the clean reads to the human reference genome using STAR software, differential analysis was performed by edgeR package. For RNA-Seq data analysis, the bulk RNA-Seq data in a form of digital gene expression (DGE) matrix was analyzed using the Seurat package for R, v. 3.1.5 [119]. Variable genes were identified using the *FindVariableFeatures* function. Top fifteen markers for each cluster were identified using a Wilcoxon Rank Sum test, and a heat map was generated using the *DoHeatmap* function.

## SUPPORTING INFORMATION

**S1 Fig. Nurr1 overexpression (OE) or knock-down (KD) substantially alters host transcriptome**. **A**, heatmaps representing top 15 gene markers for each treatment group. Statistically-significant (*p* < 0.001) differentially expressed genes were determined using the Wilcoxon rank-sum test reflecting the impacts of Nurr1 OE by comparing the control cells HC69-3X-FLAG-vector (VT) with Nurr1 overexpressing cells HC69-3X-FLAG-Nurr1 (Nurr1 OE), as well as the impacts of KD by comparing the HC69-control shRNA1 and control shRNA2 (Ctl shRNA1/2) cells with HC69-Nurr1 shRNA1 and shRNA2 (Nurr1 shRNA1/2) cells, respectively. Various cell lines were cultured in the absence (untreated) or presence of high dose (400 pg/ml) TNF-α for 24 hr. In addition, cells were given 48 hr chase after stimulation with high dose (400 pg/ml) TNF-α for 24 hr and subsequent withdrawal. **B**, heatmaps showing top 15 gene transcript markers in samples from panel **A** rearranged according to their status of treatment with TNF-α.. The most enriched gene transcripts as the result of Nurr1 overexpression or KD are listed in columns to the left. The color-coded expression pattern of each gene transcript is shown in a heatmap to the right.

**S2 Fig**. **Nurr1 overexpression mainly impacts the recovery step following TNF-α stimulation.** Trajectories of genes after stimulation with low dose (20 pg/ml) and high dose (400 pg/ml) TNF-α for 24 hr and following a 48 hr recovery (chase) after high dose TNF-α stimulation for 24 hr and subsequent withdrawal in the Nurr1 overexpression cell line HC69-3X-FLAG-Nurr1 (Nurr1 OE) were shown. Trajectories of the same genes in the control cell line HC69-3X-FLAG-vector (Ctl VT) were also shown, with a semi-transparent line connecting identical genes between the control and Nurr1-overexpressing sides of each graph. Each line represented a gene, and the Y axis values indicated the log2 expression levels. The number of genes showing each trajectory in Nurr1-overexpressing cells was shown on top. Genes that showed no change, were up regulated, and down regulated in statistically significant manner (FDR<0.05, fold change>2) were indicated with the letters n, u, and d respectively. Grouping of the different trajectories was based on gene responses during stimulation with low dose (Step 1) and high dose (Step 2) TNF-α and the recovery time after TNF-α stimulation and subsequent withdrawal (Step 3). For instance, the group of genes marked “ndu” represented genes that were not significantly changed in response to stimulation with low dose TNF-α but were down regulated with high dose TNF-α stimulation and then up regulated during the recovery (chase) period.

**S3 Fig**. **HC69-3X-FLAG-Nurr1 and HC69-3X-FLAG-vector cells strongly differ in the recovery step following TNF-α stimulation**. Genes that showed a different trajectory after TNF-α stimulation for 24 hr and following a 48 hr recovery period between HC69-3X-FLAG-vector (control) and HC69-3X-FLAG-Nurr1 cells were identified and groups containing over 100 genes were graphed. Each line represented a gene and a semi-transparent line connected identical genes between control and Nurr1-overexpressing sides of each graph. The Y axis indicated the expression level of each gene throughout the trajectory. Grouping of genes with no statistically significant changes in expression (n), up regulated (d), or down regulated (d) in the three segments was as described in **S2 Fig**.

**S4 Fig. Nurr1 overexpression (OE) substantially alters host transcriptome**. Genes involved in top differentially negatively enriched pathways in **Fig 6A** are shown in heatmaps. The values shown in the heatmap correspond to the level of differential expression between Nurr1 overexpressing cells (marked as “Nurr1”) versus vector-infected control cells (marked as “Vector”) during the chase step. The identities of the plotted pathways and genes involved in the pathways are shown on the top and to the right, respectively.

**S5 Fig. Nurr1-specific gene expression during the chase step leads to strong downregulated of genes involved in cell cycle.** Genes that exclusively change in expression during the chase step only in Nurr1 cells (see **S3 Fig**) were superimposed on the KEGG cell cycle graph. The color bar on the top right indicates the level of differential expression for each gene in Nurr1 cells during the chase step.

**S6 Fig. TNF-α stimulation leads to strong induction of NF-κB-responsive genes along with targets of multiple inflammatory cytokines.** The most enriched transcription factor binding motifs in proximity of the promoters of differentially expressed genes are shown. The size of the circles indicates the level of enrichment, while the color intensity reflects the statistical significance as shown by FDR. Positively- and negatively-enriched motifs are shown after each treatment (shown at the bottom) in the left and right panel, respectively. The identity of each motif, as annotated in the C3 lists of the MSIGDB database, is shown to the left.

**S7 Fig. Nurr1 associates with CoREST, HDAC1, G91, and EZH2 to form a transcription repression complex in microglial cells (HC69).** HC69-3X-FLAG-vector and HC69-3X-FLAG-Nurr1 cells were cultured in the absence (untreated) or presence of high dose (400 pg/ml) TNF-α for 4 hr and 24 hr respectively. A portion of these cells were also used in a chase experiment by culturing the cells for an additional 24 hr (chase) after stimulation with high dose TNF-α for 24 hr and subsequent washing with PBS (TNF-α 24h+24h). Total protein lysates from the differently treated cells were isolated and used for co-immunoprecipitation (Co-IP) with a mouse anti-FLAG monoclonal antibody. The original protein lysates (Input) and the Co-IP products were analyzed by Western blot analysis with antibodies to FLAG, CoREST, HDAC1, G9a, EZH2, and β-tubulin respectively.

## ACKNOWLEDGEMENT

This study was supported by NIH grants R01 DA043159 and R01 DA049481 to J.K. and R21-AI127252 and two Development Awards from CFAR P30-AI36219 to S.V. We thank Meenakhi Shukla for technical assistance for production of HIV-1 reporter virus. This work made use of the High Performance Computing Resource for Advanced Research Computing and flow cytometry and virology cores of the Center for AIDS Research (CFAR) at Case Western Reserve University.

## AUTHOR CONTRIBUTIONS

J.K., F.Y. and D.A. conceived of and oversaw the study. F.Y. performed all the wet bench experiments in the manuscript except as noted and along with J.K., wrote the manuscript. D.A. performed the culture of iPSC derived microglial cells and participated in data analysis and manuscript preparation. K.N. constructed the gene knock out lentiviruses and performed the ChIP-seq data analysis. S.V. processed and analyzed the RNA-Seq dataset and performed the trajectory studies and pathway analysis, participated in manuscript preparation and submitted the RNA-Seq studies performed in this project to SRA (accession number to be provided). K.L. performed the marker gene discovery for the RNA-Seq data. Y.G. performed microglial cell culture and participated in data analysis. S.S. performed the culture of iPSC derived microglial cell and participated in data analysis. All authors read the final manuscript and commented on it.

